# Molecular Dynamics studies of Long Neurotoxins against nAChR provide key insights towards design of therapeutic peptides for Dyskinesia

**DOI:** 10.1101/2023.05.22.541753

**Authors:** Shivam Pandit, Gargi Saha, Jagadeesh Kumar. D, H.G. Nagendra

## Abstract

Peptide-based drugs are widely used as therapeutic products that exhibit strong binding affinities with high specificity and low toxicity. To develop novel therapeutics against severe involuntary muscle movement and trembling disorder (Dyskinesia), the study explored the design of peptides that could target the nicotine acetylcholine receptor (nAChR) as antagonistic leads. The work aimed to suggest that the pharmacological interaction of snake venom toxins and its associated peptides with nAChR’s could be exploited as plausible solutions for PD. Molecular docking and Molecular dynamics exercises were carried out using long chain neurotoxins (α-BTx, 1NTN) to analyze the efficacies of their binding to nAChR’s, with both the entire toxin and short peptide moieties. The study demonstrated that the **novel-peptide (F8J2D7)** system appears to possess better affinity with the nAChR than the entire toxin, which is a first step towards designing peptide-based drugs from snake venom proteins that could facilitate further research in the peptide-engineering and drug designing prospects for various neurodegenerative diseases.

## 1. INTRODUCTION

Parkinson’s disease (PD) is a disorder commonly caused by the death or impairment of nerve cells, predominant in the age group of 65-69 **(Corradi & Bouzat, 2016).** Pathological studies highlight that neurons in the brain produces chemicals like Dopamine and Acetylcholine which help the brain and the nerve cells to remain active **(Dani, 2015).** In Parkinson’s disease, the nerve loses the ability to release dopamine and produces an excess amount of Acetylcholine, leading to depression, and rapid involuntary muscle movements in the legs, hands and head, which is called Dyskinesia **(Kalamida et al., 2007).** Studies showed targeting Acetylcholine receptors (AChR) exhibit a therapeutic benefit in Parkinsonism **(Quik et al., 2011).** Acetylcholine receptors are known to be major targets for various neurodegenerative diseases (NDD), like Alzheimer’s, and Huntington disease (**Ferreira-Vieira et al., 2016; Foucault-Fruchard et al., 2018**).

A subclass of AChR’s known as neuronal type nicotine acetylcholine receptor (nAChR) is the most explored one in neurodegenerative studies **(Posadas et al., 2013).** nAChR have pentameric complex structures, composed of only α (α7-α10) subunits, resulting in five identical acetylcholine binding sites. nAChR is further divided into the neuronal type and Muscle type, based on the specific combinations of subunits, mediating different physiological roles **(Carlson & Kraus, 2021; Pedersen et al., 2018).**

Neuronal nAChR forms a family of subunits composed of α2-α10 and β2-β4 subunits. In mammalian neurons, eight α-subunits (α2-α7, α9, and α10) and three β-subunits (β2-β4) are identified. The extracellular domain of α subunits forms homomeric and heteromeric pentamers that contain cysteine groups **(Kruse et al., 2014).** The hetero-pentamers α4β2, and homo-pentamers of α7 and α9 show high-affinity for nicotine and acetylcholine ligands due to the presence of cysteine residues **(Bohnen & Albin, 2011).** Similarly, Muscle-type-nAChR is made up of α1, β1, γ, and δ/ε (adult) subunits in a 2:1:1:1 ratio that receives acetylcholine for muscular contraction. The muscle-type nAChR are assembled via different subunits forming various arrangement (**Kruse et al., 2014; Pedersen et al., 2018**). Muscarinic Acetylcholine receptors (mAChR) are G protein-coupled receptors (GPCR) and are categorized into 5 subfamilies viz. M1 to M5 encoded by the genes CHRM1 to CHRM5 respectively, which plays a central role in human physiology, regulating heart rate, smooth muscle contraction, glandular secretion and many fundamental functions of the central nervous system (CNS). Three of those receptor subtypes (M1, M3 and M5) exhibits simulated regulation whereas the remaining two subtypes (M2 and M4) exhibits inhibitory regulation (**Giastas et al., 2018; Haga, 2013; Svoboda et al., 2017**).

Studies showed nAChR and their subtypes regulate the cationic channel within the neuronal pathway **(Posadas et al., 2013).** They are ubiquitously distributed in peripheral and central nervous systems playing a role in postsynaptic activity, exciting the neurons, or as presynaptic receptors, modulating the release of many neurotransmitters (**Dani, 2015; Kalamida et al., 2007**). Recent studies have suggested that targeting nAChR may be a useful strategy to improve conditions in Parkinson’s disease **(Scarr, 2012).**

It is found that snakebite injuries are considered a major socio-medical problem in the underdeveloped and developing parts of the world, as a large number of people suffer and die because of snake venom poisoning. Out of 2.7 million snake bitten population approximately 400,000 survivors suffer from permanent physical disabilities and around 81,000–138,000 victims die **(Girish et al., 2019).** In hundreds of microliter of snake venom, a cocktail of several low molecular weight compounds and peptides/proteins that cause local and systemic malfunctions of normal cells and organs by targeting specific receptors, ion channels, with selectivity and affinity **(Girish et al., 2019).** Despite snake venom being highly toxic and poisonous, research studies suggest that, their proteins possess pharmacological effects like hemotoxic, neurotoxic, cytotoxic and proteolytic properties **(Munawar et al., 2018)**. They also exhibit potential therapeutic values as a neuroprotective and neurodegenerative agents **(Gazerani, 2021).** This point could be exploited towards providing therapeutic effects for dyskinesia **(Gazerani, 2021),** since snake venom proteins bind to the Ligand Binding Domain (LBD) of nAChR’s resulting in the blocking of the acetylcholine (Ach) binding site **(Young et al., 2003).**

Literature studies and databases identified approximately 59 important protein families with their subtypes, which are categorized based on the quantity and availability of proteins present in the pool of venom **(Tasoulis & Isbister, 2017).** Snake venom protein families are divided into a) Four dominant family groups namely PLA2, SVMP, SVSP and 3FTx b) Six secondary protein families namely KUN, CRiSP, LAAO, CTL, DIS, and NP c) 9 minor protein families, like acetylcholinesterase, hyaluronidase, nucleotidases, phosphodiesterase, phospholipase B, nerve growth factor, vascular endothelial growth factor, and snake venom metalloproteinase inhibitor d) 36 rare protein families e) four “unique” protein families, which are restricted to a single genus (**Tan et al., 2016; Tasoulis & Isbister, 2017**).

Out of these 59 protein families, three-finger toxins (3FTx) is one of the largest family of snake venom proteins with good evidence of therapeutic properties (**Tan et al., 2016; Tasoulis & Isbister, 2017**). α-neurotoxins compete with Acetylcholine (Ach) and bind to AChR’s as antagonists by blocking the action of ACh to the postsynaptic membrane, thus inhibiting the ion flow and leading to neuronal death **(Nirthanan & Gwee, 2004).** AChR blockers or Acetylcholine antagonists can be used to prevent the chemical reaction and maintain a balance between the neurotransmitters **(Posadas et al., 2013).** For example, alpha-bungarotoxin (α-BTx), a long neurotoxin from ***Bungarusmulticinctus*** has shown therapeutic effects against neurodegenerative diseases (**Nirthanan & Gwee, 2004; Young et al., 2003**).

Hence this particular study aims to understand the nature of interaction of long neurotoxin derived from Najaoxiana (Central Asian cobra) (Oxus cobra) with neuronal type homomeric α7 nAChR, so as to explore the design of pharmacophorically derived peptides, as a probable cure for dyskinesia.

## 2. MATERIALS AND METHODS

### 2.1 Sequence and Structural data collection and analysis

Snake venom long neurotoxins from different snake families were retrieved from Uniprot database **(Bateman et al., 2021),** and are listed in Table 1. Global pairwise sequence alignment and Multiple sequence alignment (MSA) studies were performed for these retrieved sequences against the standard long chain neurotoxin i.e. alpha-Bungarotoxin (Uniprot ID -P60615 and PDB ID-1IK8) from ***Bungarismultocinctus*** using EMBL-EBI pairwise sequence alignment and ClustalW tools **(Needleman & Wunsch, 1970; Sievers et al., 2011**), and is depicted in Figure 1. Sequence identities and similarity percentages were calculated for all the 25 retrieved sequences. It was found that the long chain neurotoxins Alpha-elapitoxin-Nno2 (Alpha-EPTX-Nno2a) derived from ***Najaoxiana* (Central Asian cobra) (Oxus cobra)** with Uniprot ID-P01382 and PDB ID-1NTN, showed the highest identity and similarity percentage w.r.t to the standard i.e. alpha-Bungarotoxin (PDB ID-1IK8), as indicated in Table 1.

**Table. 1:**
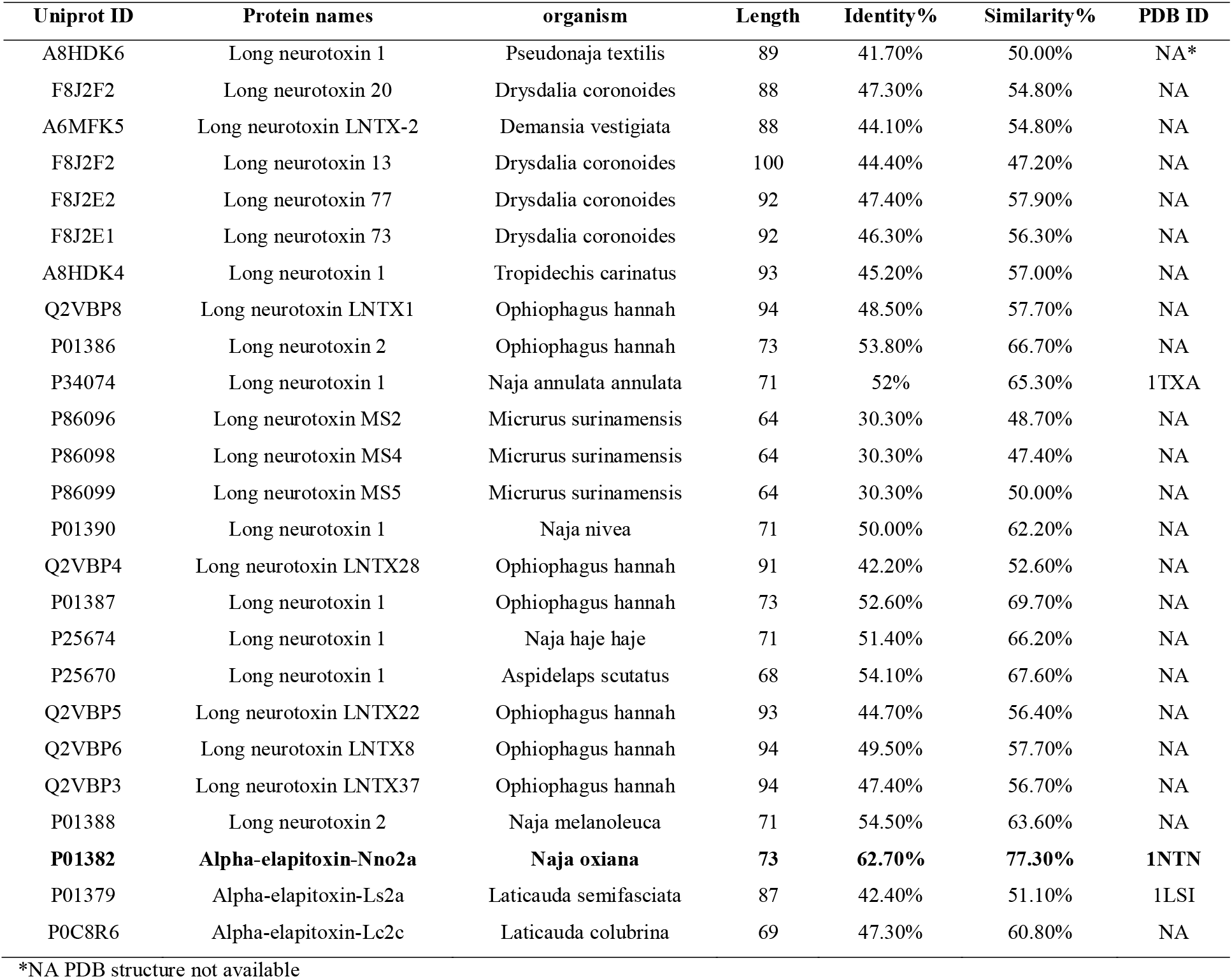
List of long neurotoxins retrieved from Uniprot database. The highlighted Uniprot id is having highest Identity % and Similarity % with respect to α-BTx (PDB ID-1IK8).

**Figure 1:**
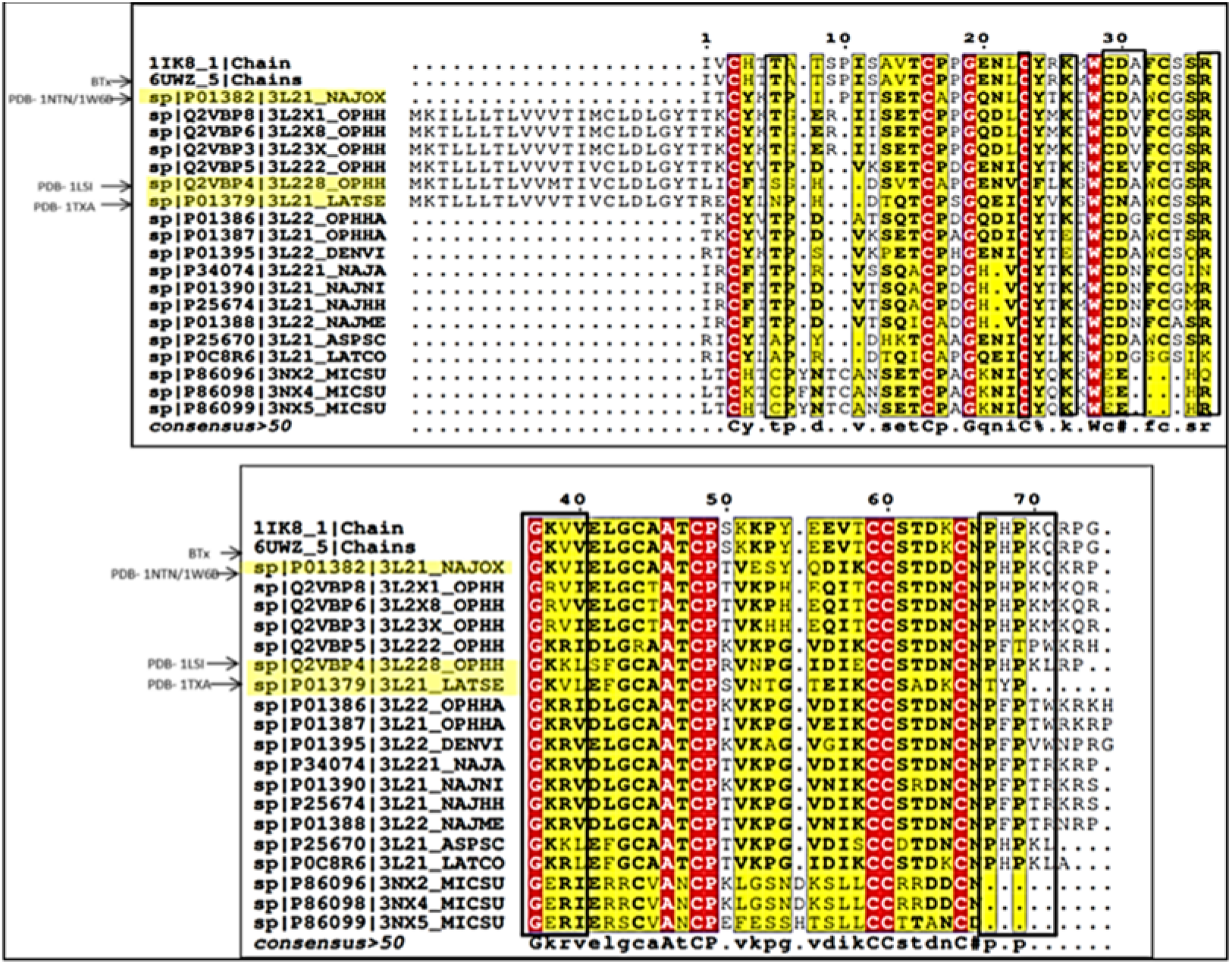
Multiple sequence alignment (MSA) of all long neurotoxins keeping α-BTx as standard.

The corresponding three-dimensional structures of the free forms of snake venom long chain neurotoxin (1NTN, 1IK8), and the complexed structure of α-BTx long chain toxin with nAChR (PDB ID - 4HQP), were retrieved from the Protein Data Bank (PDB**) (Berman et al., 2000).** The structural similarity exercise were carried out by superimposing the α-BTx bound complexed structure (4HQP) against the α-BTx unbound form (1IK8) and the long chain neurotoxin from Alpha-EPTX-Nno2a (1NTN) (unbound state). The RMSD value for the C-alpha atoms, backbone atoms, side chain atoms, and all the atoms, were calculated and are listed as in Table 2. Analyzing the RMSD values over the bound and unbound states of toxins, the protein modeling studies for 1NTN neurotoxin sequence was conducted using 4HQP as a template via SWISS Model server, which is a web-based integrated protein homology modelling server responsible for building protein homology models at different levels of complexity **(Waterhouse et al., 2018).** Through this exercise, the structural conformation of 1NTN long chain toxin was brought to close proximity with the experimental model α-BTx (4HQP). The RMSD values of these remodeled structure, were also re-estimated (Table 2). This 1NTN remodeled structure has been used for the further investigations related to molecular level interactions, peptide modeling and dynamics.

**Table 2:**
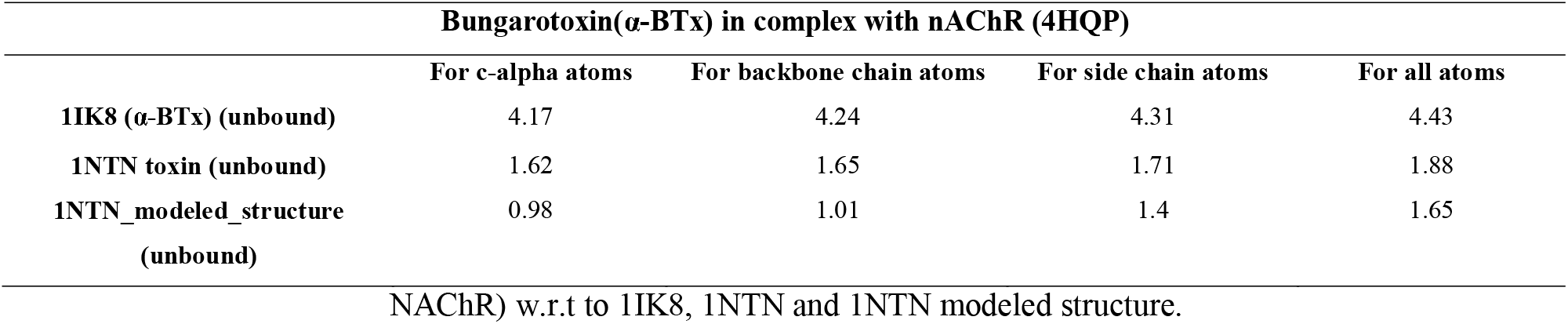
RMSD values in Angstroms between the long chain neurotoxin structures 4HQP (α-BTx in complex with

### 2.2 Molecular Docking Studies

Docking studies for the long chain toxins (α-BTx, 1NTN) and the receptor of interest (nAChR) were conducted via the online web based server HDOCK **(Yan et al., 2020),** which is a novel protein-protein docking platform that uses hybrid docking algorithm, associated with fast Fourier transform (FFT)–based global search and iterative knowledge-based scoring function. 100 docked poses were generated for each complex, and among them the best pose based on the interacting residues and binding energies scores were carefully examined. The residue level interactions (within 4.5 Angstroms) were also tabulated using Discovery Studio visualization platform **(Dassault Systèmes, 2020)** for the best pose that had the minimum binding energy (Table 4).

**Table 3:** probable mutations occurred across the various position of the peptide stretch obtained from MSA study; important positions are highlighted.

| Position | α-BTx peptide | 1NTN_peptide | Probable Mutations |
| --- | --- | --- | --- |
| 1 | R | T | M,R,L,K |
| 2 | K | K | R,E |
| 3 | M | T | M,K,S,V,A |
| 4 | W | W | Conserved |
| 5 | C | C | Conserved |
| 6 | D | D | A |
| 7 | V | A | I,R,D,V,N,G,F,Y |
| 8 | F | W | F,L,S,R |
| 9 | C | C | Conserved |
| 10 | S | G | F,S,T,L,A,H |
| 11 | S | S | I,D,T,M,Q,N,F,V |
| 12 | R | R | K,N,L |
| 13 | G | G | L |
| 14 | K | K | R,E |
| 15 | V | V | R,K,I |
| 16 | V | I | V,L,E,M |
| 17 | E | E | D,V,N,S,Q |
| 18 | L | L | M,F |
| 19 | G | G | Conserved |
| 20 | C | C | Conserved |

**Table 4:**
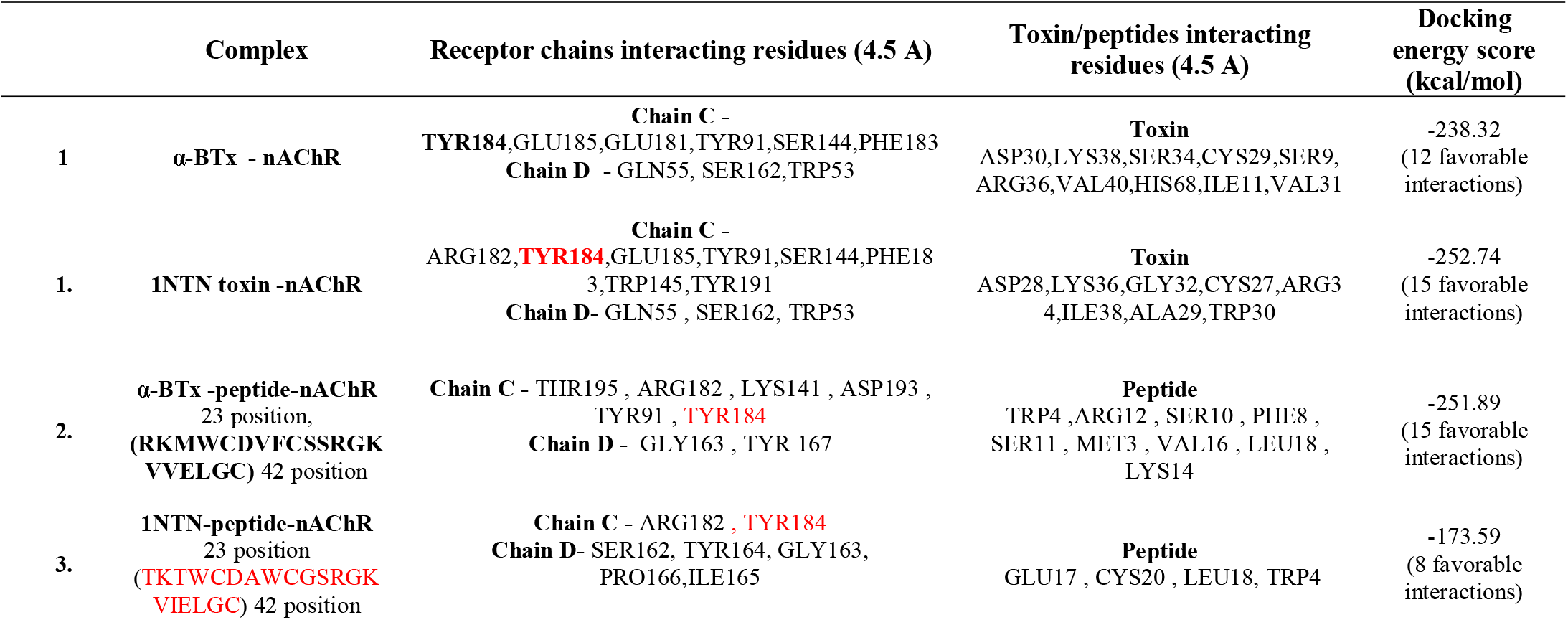
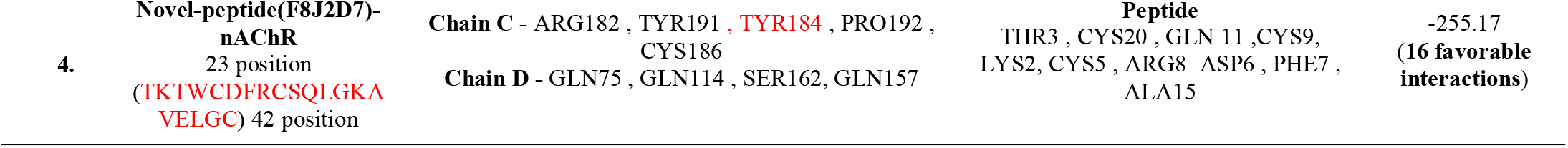
Details of Interacting residues details of the receptor, toxin and peptide obtained from the best docked poses.

### 2.3 Peptide Modelling Studies

Analyzing the residue level binding chemistry from the receptor-toxin complexes, we noticed that, a stretch of 20 amino acids from long chain toxins showed maximum interaction with the nAChR pocket residues. Hence, in order to explore the possibility of a short peptide that would exhibit similar binding properties, the modeling exercise using denovo peptide structure prediction web server PEP-FOLD3 **(Lamiable et al., 2016),** was carried out. PEP-FOLD3 is a novel computational framework, which allows both (i) *de novo* free or biased prediction for linear peptides between 5 and 50 amino acids, and (ii) the generation of native-like conformations of peptides interacting with a protein when the interaction site is known in advance. Among the 10 conformations generated, the energetically most stable peptide conformation was used for docking studies with nAChR and the associated binding scores were estimated. Further, the residue level binding interactions (within 4.5 Angstroms) were tabulated as well, and compared with the earlier values for the overall toxin structure.

### 2.4 Identification of novel peptide sequence from multiple sequence alignment (MSA) studies

To further explore the possibility of identifying any novel peptide, that could possibly exhibit the best binding property with the receptor, a detailed sequence analysis was conducted from the family of long chain 1NTN neurotoxins. BLAST **(Altschul et al., 1990)** exercise was performed and about 112 homologous sequences were retrieved. Multiple sequence alignment exercise was conducted for these 112 sequences (Figure 4a), and the probable mutations at various positions across the 20 amino acid stretch were carefully listed (Table 3). During this exercise we noticed that, a novel sequence with **UNIPROT ID – F8J2D7, which is a long neurotoxin 43**, from ***Drysdalia coronoides (White-lipped snake)***, possessed a sequence identity of 65% and similarity of 70% w.r.t to standard 1NTN peptide stretch (Figure 4b). The mutational mapping exercise between F8J2D7 and 1NTN peptide stretch suggested interesting mutations at the 7^th^ position and 8^th^ position of the sequence. This novel 20 aa sequence was also used for a detailed study, to ascertain its efficacious binding properties.

**Figure 4a:**
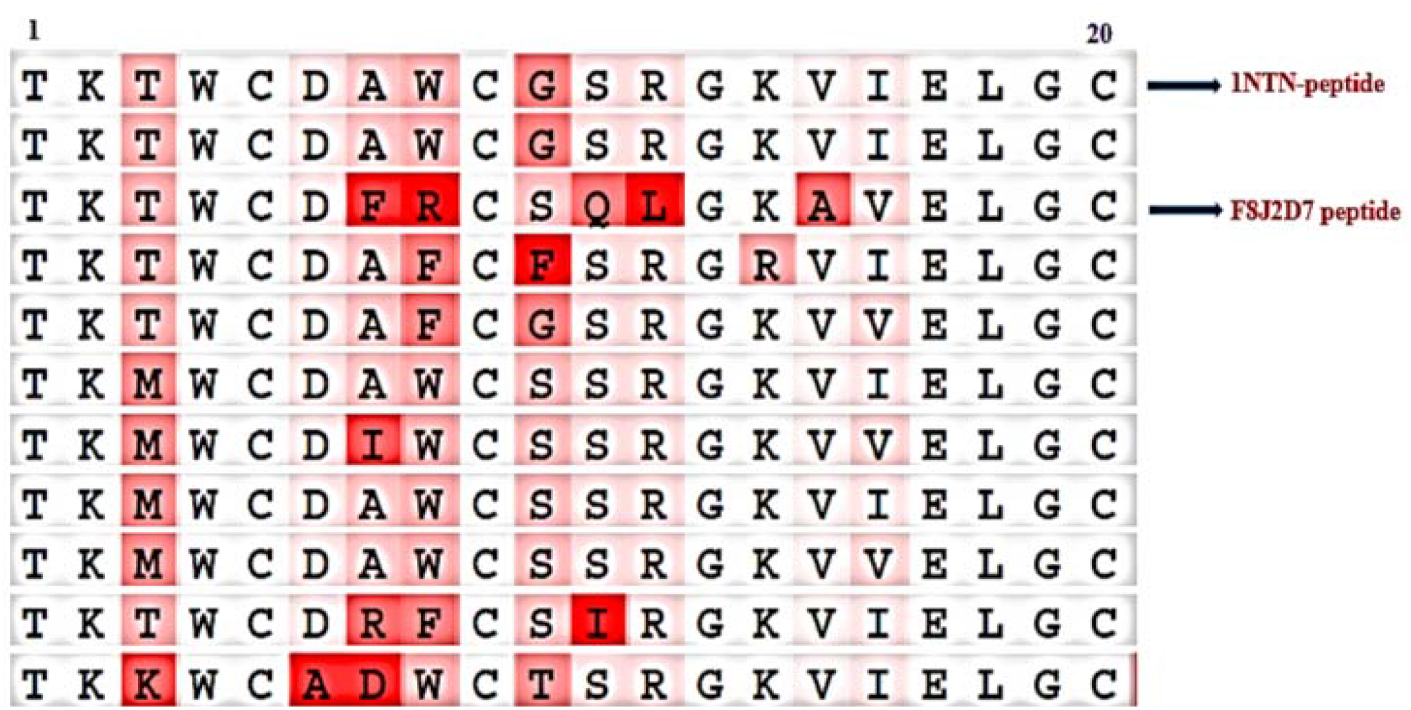
Multiple sequence alignment of 1NTN peptide stretch

**Figure 4b:**
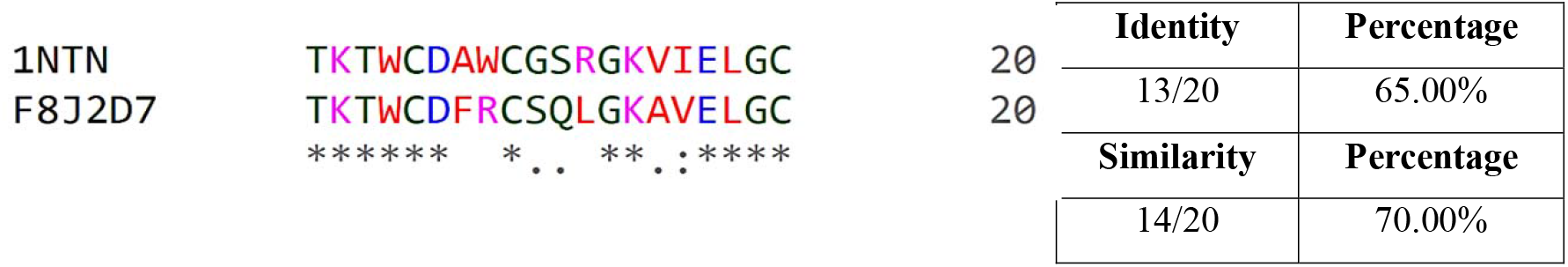
Sequence alignment of 1NTN-peptide and novel peptide

### 2.5 Molecular Dynamics simulation studies

To understand the stability of docking interactions, Molecular Dynamics studies for the best receptor toxin and receptor –peptide complexes were carried out using GROMACS 2020 software package **(Bekker et al., 1993).** The Protein parameters for the simulation runs were generated using OPLS-AA/L all-atom force field (2001 amino acid dihedrals). Before the production run, the complex structures were solvated inside a cubic box of the transferable intermolecular potential TIP3P water model **(Jorgensen et al., 1998).** Under the periodic boundary conditions, the residues are set according to the standard ionization states at pH 7.0. The entire complex was neutralized using NaCl ions which were added through Monte-Carlo ion-placing method. The simulation was conducted in three stages and 1000 kJ/mol.nm2 force constant was used for restraining all heavy atoms preserving original protein folding. First stage deals with the initial optimization of each system geometry for 5ps. Following this, a two-staged equilibrium, where the whole system was trained for 100,00 iterations (100 ps) at each stage, was performed. The first equilibration was done under a constant number of particles, volume, and temperature (NVT) ensemble concerning the Berendsen temperature coupling method for regulating the temperature within the box **(Hess et al., 2008).** Second equilibration was done under constant (NPT) ensemble at 1 atm and 303.15K using the Parrinello-Rahman barostat **(Hess et al., 2008).**

The Dynamics production run for the complexes of interest were carried out for 70ns under constant pressure. Furthermore, the simulation trajectories were used to calculate the root mean square deviation (RMSD), root mean square fluctuation (RMSF), radius of gyration (ROG) values using gmx RMSD, gmx RMSF and gmx gyrate tools respectively **(Bekker et al., 1993).** The graphical plot were generated using the grace package **(Turner et al., 2005),** as depicted in (Figure’s 6a-6e, 7a-7e, 8a-8e).

### 2.6 Average binding free energy calculation (MM/PBSA)

MD-based Molecular Mechanics/ Poisson Boltzmann Surface Area (MM/PBSA) approach was used to calculate the average binding free energy of the complexes of interest **(Baker et al., 2001; Kumari et al., 2014).**This tool help us to evaluate the binding free energy of the systems. A total of 100 frames at an interval of 400ps difference were extracted from the last 30-70ns (40 ns) stable trajectory, and the average total binding energy of the complexed system was calculated. Further, the MMPBSA energy components such as electrostatic, van der waals, polar solvation energy, and non-polar solvation energy were also calculated and tabulated (Table 8). The net energy of the system was estimated through the following equation, Δ**G _Binding_ =**Δ**G _Complex_ -** Δ**G _Receptor_ -** Δ**G _Inhibitor._**

## 3. Results and Discussion

### 3.1 Sequence and Structural analysis

Fasta sequences of 20 long chain toxins from different organisms belonging to three finger toxin family (3FTx), were retrieved from the Uniprot database, and a sequence comparison study, with the standard alpha-Bungarotoxin (α-BTx), was conducted. Interestingly, the alignment showed that, these sequences had segments of conserved residues across the stretch. In total, 7 cysteines, 2 Glycine’s, 1 tryptophan, 1 alanine and 1 proline residues, were found to be conserved across the retrieved sequences as in (Figure.1) (highlighted). Amongst the conserved residues, the tryptophan (28^th^ position) and the glycine residues (37^th^ position) were found to be a part of the interaction site of toxins with the nAChR pocket. On the other hand, the conserved cysteines were responsible for disulfide bonds, and the proline and alanine residues were part of the loops in the toxin structure.

The alignment results, exhibited that amongst the pool of neurotoxins the Alpha-elapitoxi-No2a (Alpha-E Najaoxiana (Central Asian cobra - Oxus cobra) long chain toxin with Uniprot id P01382 (1NTN) had the highest percentage similarity of 77.30% and an identity percentage of 62.70% as indicated in Table 1 and highlighted Figure 2, with the standard α-BTx (1IK8).

**Figure 2:**
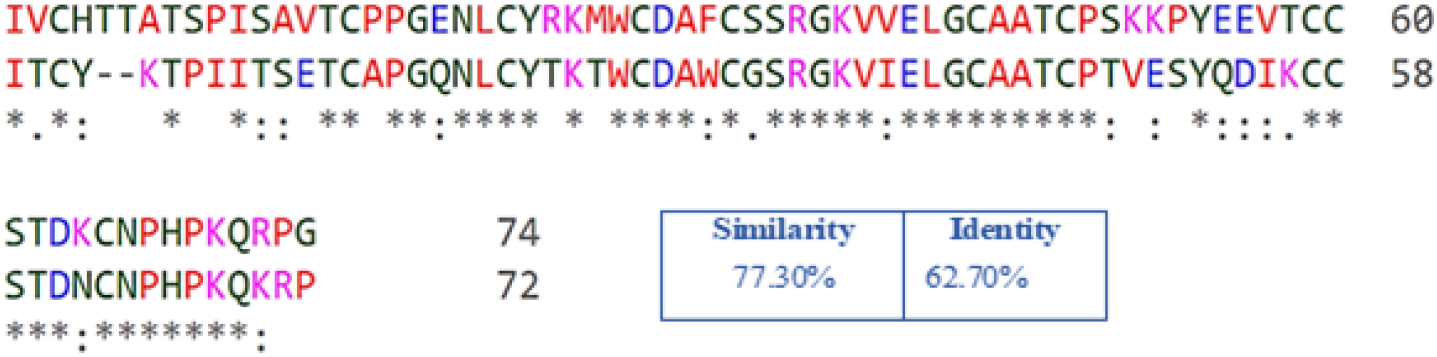
Sequence alignment of 1IK8 and 1NTN

Further, to understand the structural orientations of the (unbound state) of long chain neurotoxins (1IK8, 1NTN) w.r.t to the ligand bound experimental structure (4HQP), a structural level analysis was carried out by superimposing the toxins of interest. The RMSD (root mean square deviation) values for 1NTN, α-BTx (1IK8), keeping α-BTx complex (4HQP) as a reference, were calculated, for all the c-alpha atoms, backbone chain atoms, side chain atoms and all atoms respectively (Table 2).

From the superimposition analysis, we noticed that, both the unbound toxins i.e., 1IK8 and 1NTN showed structural deviations w.r.t to the reference structure (4HQP). The 1IK8 free structure was found to have a higher RMSD profile than 1NTN, indicating that, the movement of atoms occurs in the bound stable state compared to an unbound free structure (Table 2). Template based modeling studies were carried out using 4HQP as a template for long chain toxin (1NTN) (unbound state) and the structural superimposition and RMSD calculation exercise were again carried out for the remodeled structure. The results exhibited that, the remodeled structure had a closer RMSD profile w.r.t to standard 4HQP (Table 2), suggesting that, both the structures i.e., the ^α^-BTx complex (4HQP) and the remodeled structure of 1NTN were in close conformations. This remodeled structure of 1NTN was used for all further analysis.

### 3.2 Molecular Docking Studies

S. Huang et al., (2013), has observed that α-BTx (long chain neurotoxin) binds to neuronal type homopentameric α7 nAChR with a high affinity. This crystal structure of α-BTx, in complex with α7 nAChR (4HQP) was retrieved from PDB and the ligand molecule (α-BTx) was re-docked to the receptor (nAChR) using Hdock server. The interacting residues from the receptor i.e. chain C and Chain D and the residues from the α-BTx (chain F) were delineated within a range of 4.5 angstroms and compared with interactions mentioned in the research article related to the experimental structure (**Huang et al., 2013**). It was ascertained that **TYR184** from the chain C loop of the receptor is a key amino acid having critical roles in the physiological process. Based on this, site-specific docking was carried out for long chain toxin (1NTN) with the receptor (nAChR) using HDOCK server, by defining the key interacting residues from chain C loop of nAChR (**TRY184, PHE183, ARG182, GLU185**) within a distance constraint of 4.5 angstroms. The details were tabulated in the Table 4.

From the docking interaction analysis, it was noted that, 1NTN-nAChR complex had a maximum number of **15 favorable interactions** within 4.5 angstroms (associated with 11 residues from receptor chain and 8 residues from toxin chain) compared to our standard α-BTx-nAChR complex, which exhibits **12 favorable interactions** (9 residues from receptor chain and 10 residues from toxin chain). The 1NTN-nAChR complex showed 6 hydrogen bonds at chain C via the residues **TYR184, GLU185, TYR91, SER144, PHE183**, along with 4 hydrophobic interactions from **TRP145, TYR91, TYR184, and TYR191** and 1 electrostatic interaction due to **ARG182**. Further, 3 hydrogen bonds were noted with the chain D via the residues **GLN55, SER162 and TRP53** and 1 hydrophobic interaction found at **TRP53** (Table 4). In case of α-BTx-nAChR complex, we noticed that, it had 7 hydrogen bonds at chain C from **TYR184, GLU185, GLU181, TYR91, SER144, PHE 183**, along with 2 hydrophobic interaction via **PHE183 and TYR184** respectively. At chain D, 2 hydrogen bonds were noted due to **GLN55 and SER162,** along with 1 hydrophobic interaction via **TRP53** residue (refer Table 4). Additionally, the associated docking energy scores for the respective complexes was found to be **-252.74 kcal/mol** (1NTN-nAChR complex) compared to **-238.32 kcal/mol** for the α-BTx-nAChR complex respectively as indicated in Table 4.

Moreover, the binding conformation analysis of these docked toxins around the binding pocket of the receptor (nAChR), revealed that both the long chain toxins (α-BTx and 1NTN) had a similar nature of binding with favorable number of interactions with nAChR pocket residues (Figure 3, 5a-5d). Thus, the bound toxin molecules was used as reference, for further peptide modelling, binding and Molecular dynamics studies.

**Figure 3:**
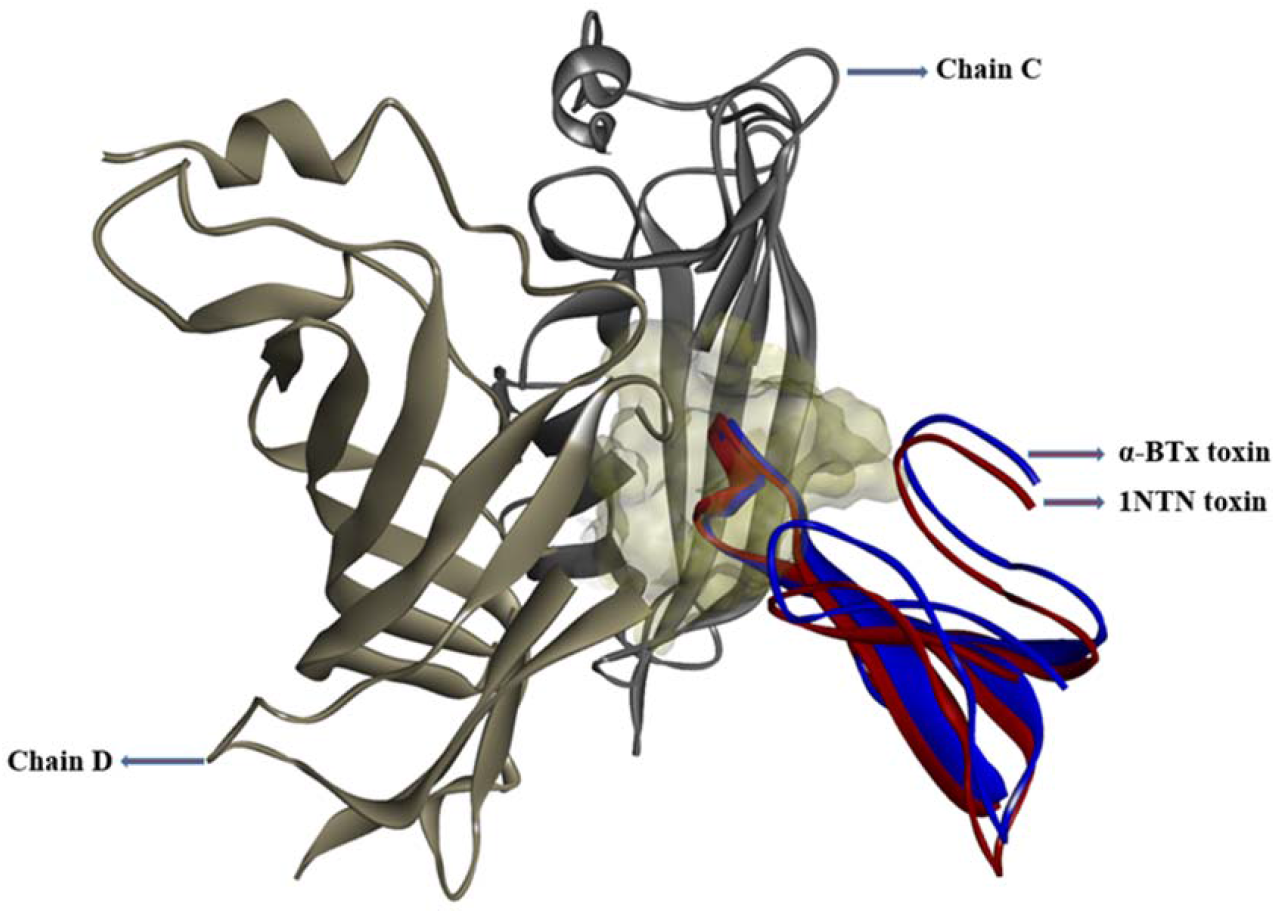
Nature of binding of α-BTx and 1NTN long neurotoxin inside the binding pocket area of nAChR

### 3.3 Studies with peptides

From the residual level interaction chemistry with the receptor (nAChR) and the toxins (α-BTx and 1NTN), it was delineated that a stretch of about 20 amino acids from long chain toxins (α-BTx and 1NTN), showed maximum interaction with nAChR pocket residues, which could be explored for designing peptide, that could exhibit similar nature of binding chemistry with the receptor (nAChR) and provide similar pharmacological effect as the entire toxin molecules.

Along with the peptide designing exercise, we further wanted to use the 20 amino acid stretch from1NTN long chain toxin, and derive a novel peptide sequence from the pool of similar sequences. Towards this, a total of 112 peptide sequences were retrieved from BLAST and used for sequence alignment study (Figure 4a) (supplementary material). A mutational mapping exercise was carried out to fish out all the probable mutations across the 20 amino acid stretch (Table 3). Interestingly, from the alignment results, we found that, one novel sequence with **UNIPROT ID – F8J2D7 (long neurotoxin 43)** from ***Drysdalia coronoides (White-lipped snake)***, with a sequence identity of 65% and similarity of 70% w.r.t to standard 1NTN stretch (Figure 4b) exhibited interesting mutations at multiple positions across the sequence stretch (Table 3). The mutation at the 7^th^ position was ALA>PHE (A>F) residue, which is from non - polar small hydrophobic residue to a large aromatic, hydrophobic amino acid residue. Likewise, at the 8^th^ position the mutation was from TRP>ARG (W>R) residue; which is from an non-polar aromatic hydrophobic residue to a large positively charged polar residue (Figure 4a). After a detailed investigation, of 1NTN-nAChR complex we found that, these positions were crucial and had interactions with the critical residues of chain C loop of nAChR (**TYR184, ARG 182, PHE183**). The mutations at these critical positions for **the novel sequence** were found to be interesting, and hence, the consequences of these mutations with respect to its conformation and binding affinities were analyzed in detail.

The 20 amino acid stretches from α-BTx, 1NTN and the novel sequence **F8J2D7**, were used for peptide modeling exercises (via PEPFOLD3 sever) and the energetically best modeled structure was analyzed in each case. Further, docking studies with nAChR was carried out in site specific manner using HDOCK server for these modeled peptides, and the best poses were evaluated based on the number of interacting residues, binding energy scores and binding conformations respectively (Table 4).

The analysis, suggested that both the α**-BTx-peptide-nAChR complex** (with **15 favorable interactions** across 8 residues from receptor chain and 9 residues from peptide moiety) and the **novel-peptide (F8J2D7)-nAChR complex (with 16 favorable interactions** across 9 residues from receptor chain and 10 residues from peptide moiety) showed similar number of favorable interactions within 4.5 angstroms with the nAChR pocket residues (Table 4) (Figure 5e-5j) .On the other hand, in case of **1NTN-peptide-nAChR complex** (7 residues from receptor chain and 4 residues from peptide moiety) lesser number of favorable interactions **(8 favorable interactions)** with the nAChR pocket residues (refer Table. 4) (Figure 5g-5h), indicating lesser affinity of this peptide than the other two peptide systems.

**Figure 5a:**
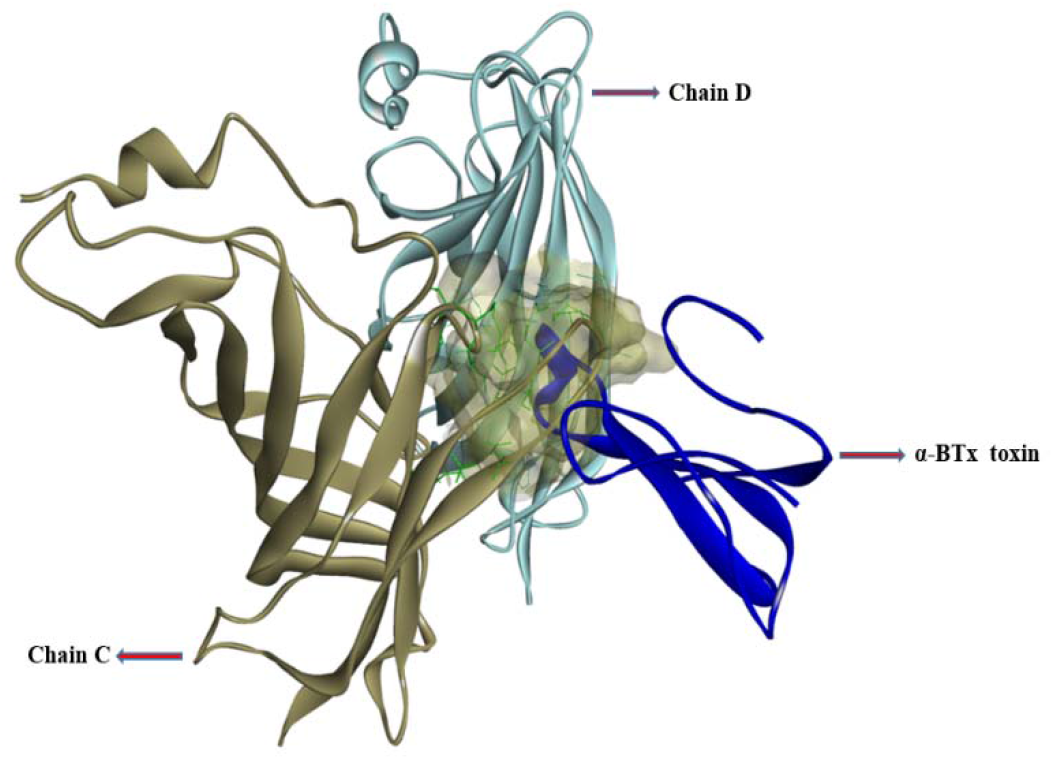
Binding nature of α-BTx toxin inside the binding pocket area of nAChR;

**Figure 5b:**
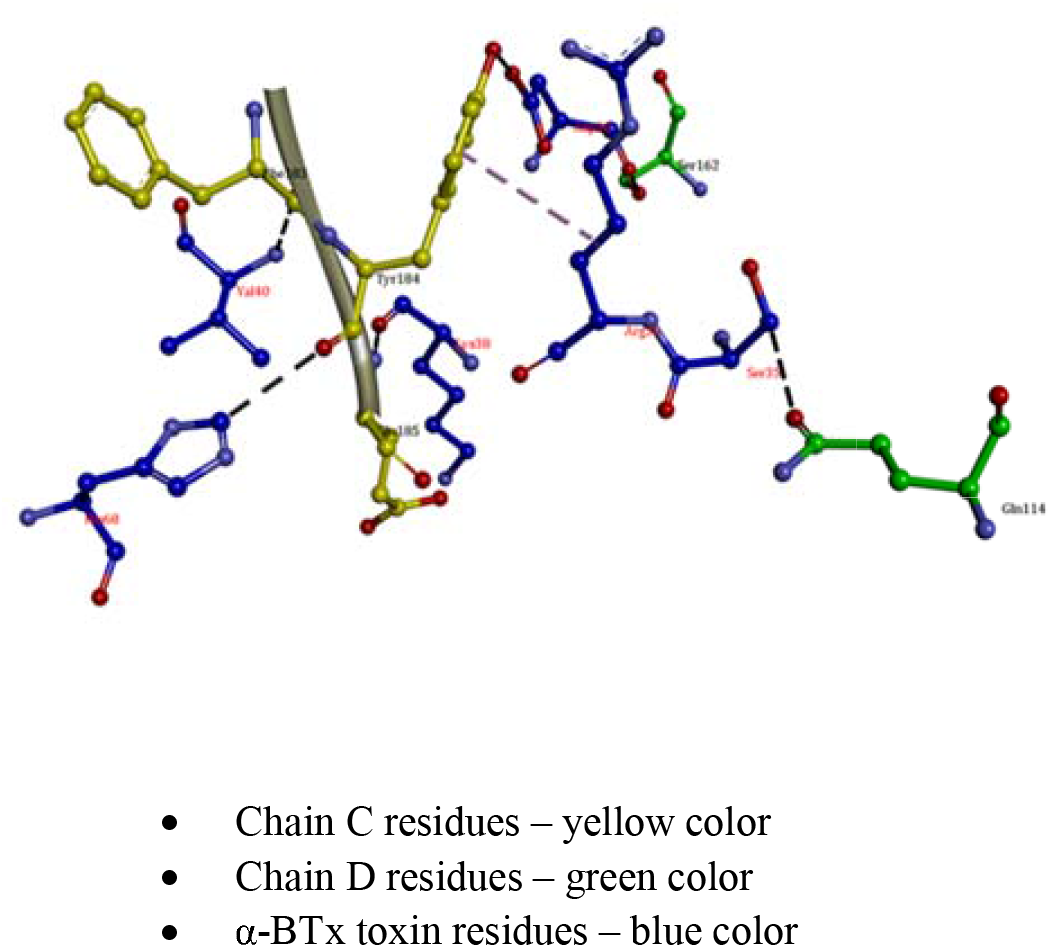
3D interaction details of α-BTx toxin with nAChR pocket residues

**Figure 5c:**
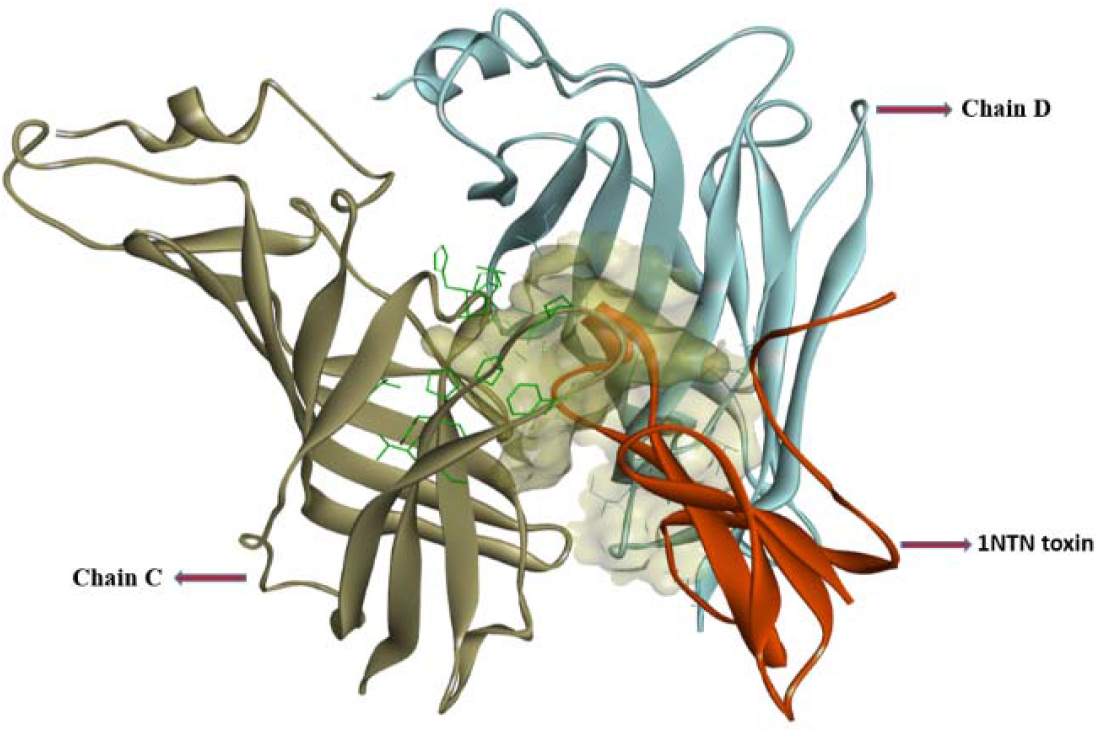
Binding nature of 1NTN-toxin inside the binding pocket area of nAChR;

**Figure 5d:**
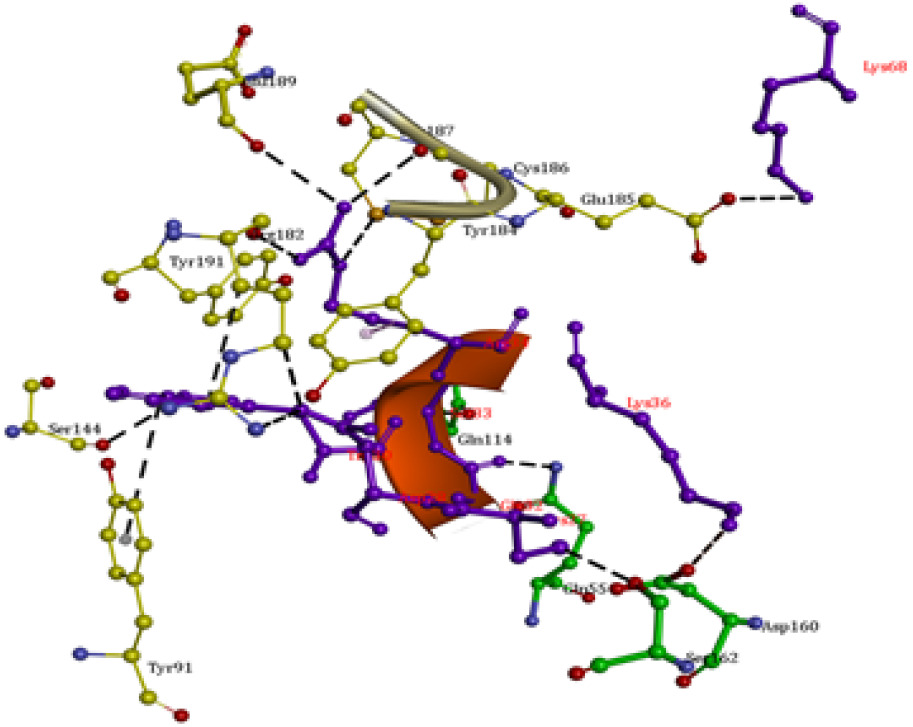
3D interaction details of 1NTN - toxin with nAChR pocket residues.

**Figure 5e:**
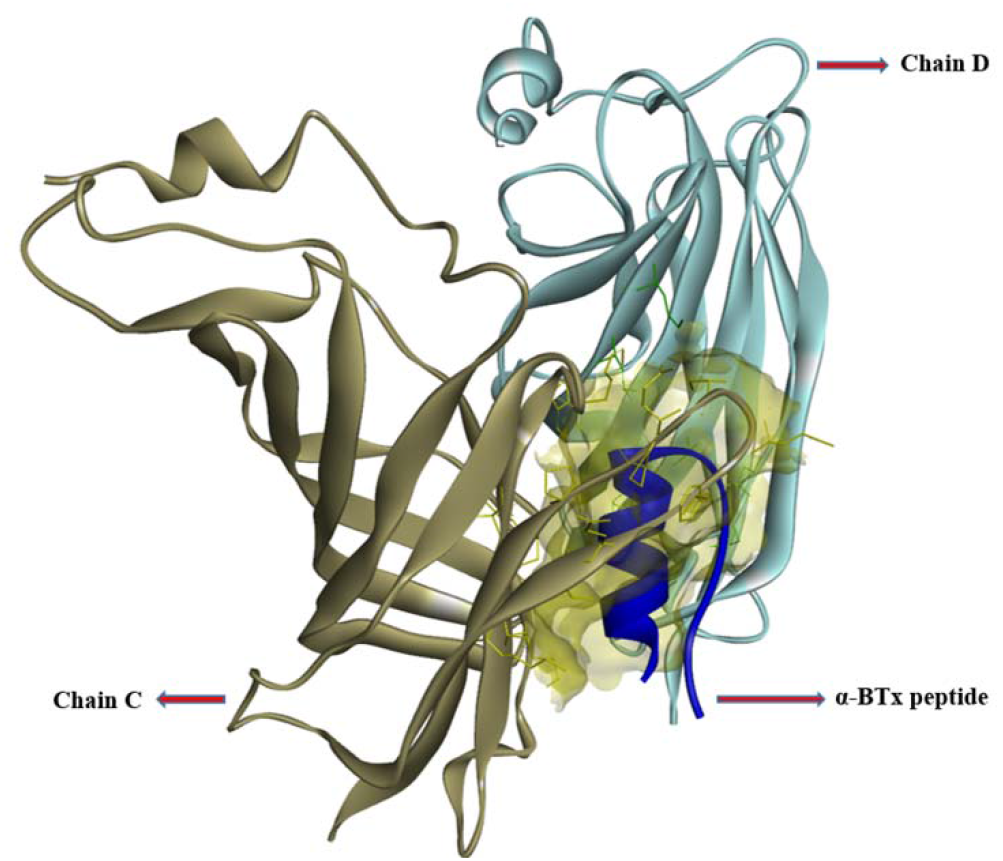
Binding nature of α-BTx peptide inside the binding pocket area of nAChR;

**Figure 5f:**
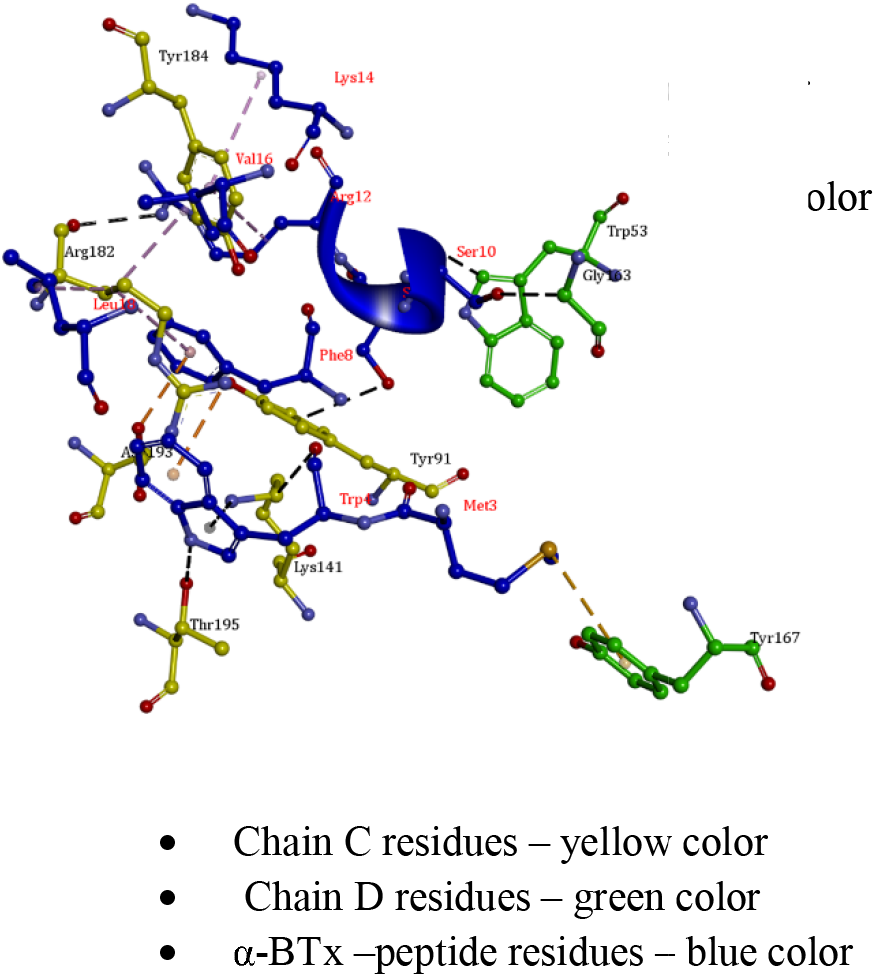
3D interaction details of α-BTx -peptide with nAChR pocket residues.

**Figure 5g:**
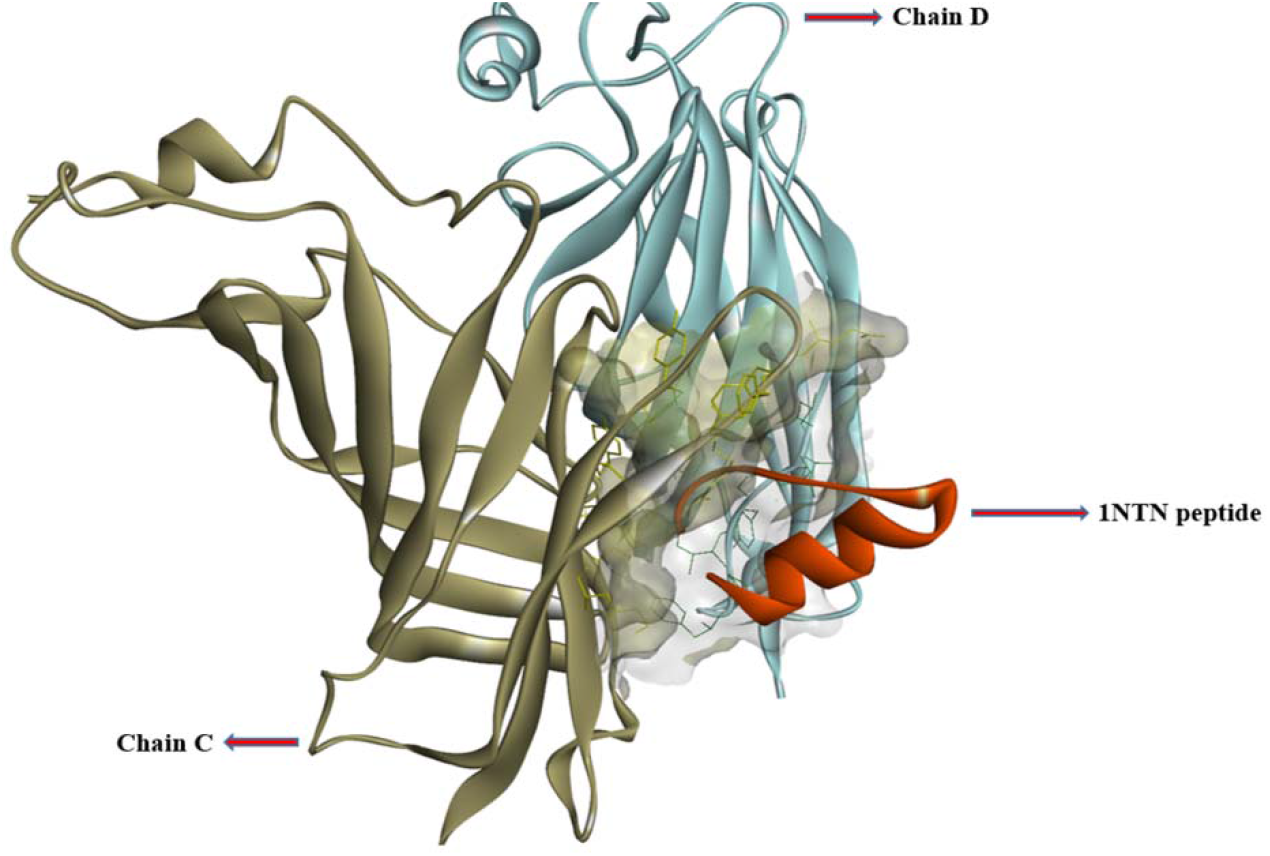
Binding nature of 1NTN-peptide inside the binding pocket area of nAChR;

**Figure 5h:**
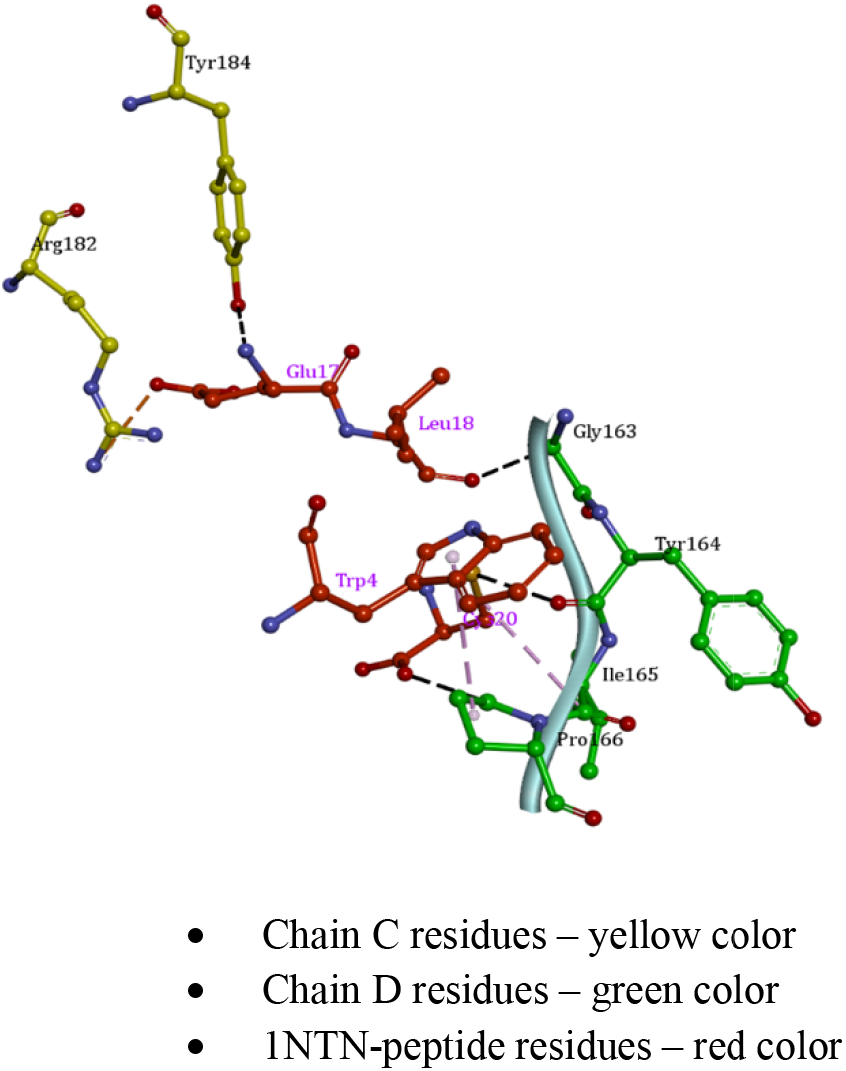
3D interaction details of 1NTN-peptide with nAChR pocket residues

**Figure 5i:**
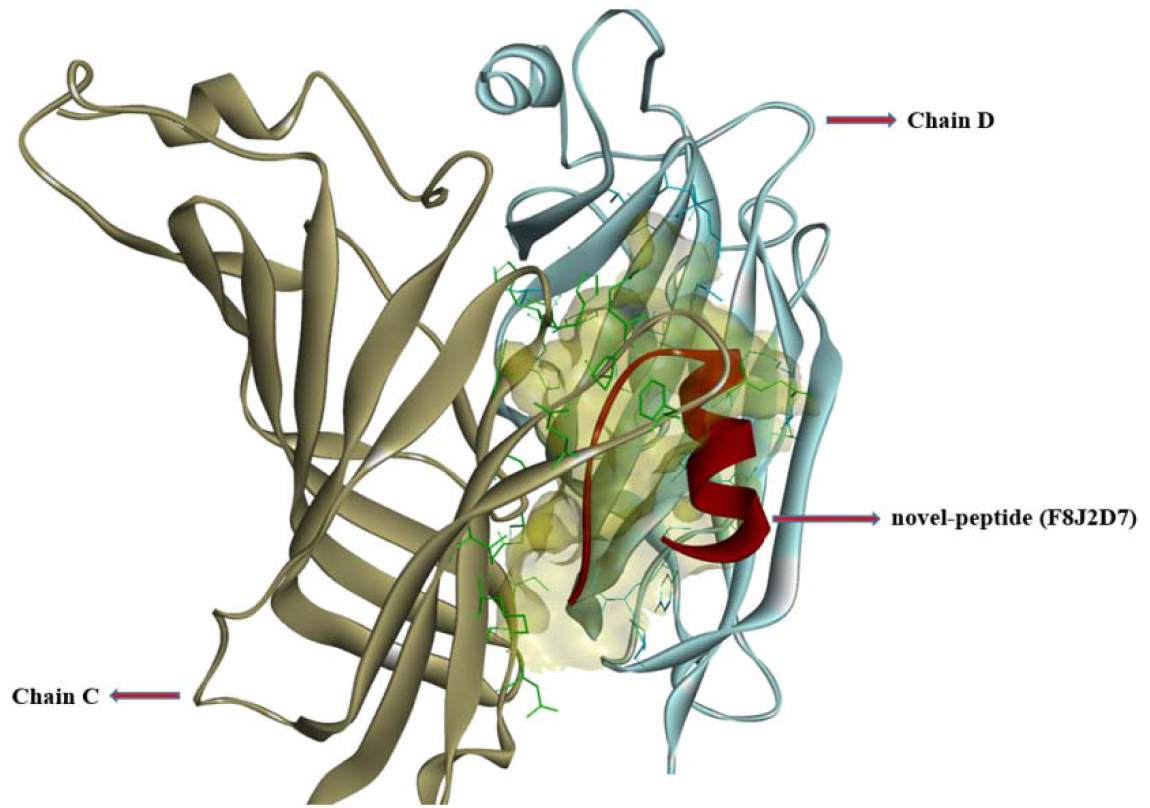
Binding nature of novel (F8J2D7) peptide inside the binding pocket area of nAChR;

**Figure 5j:**
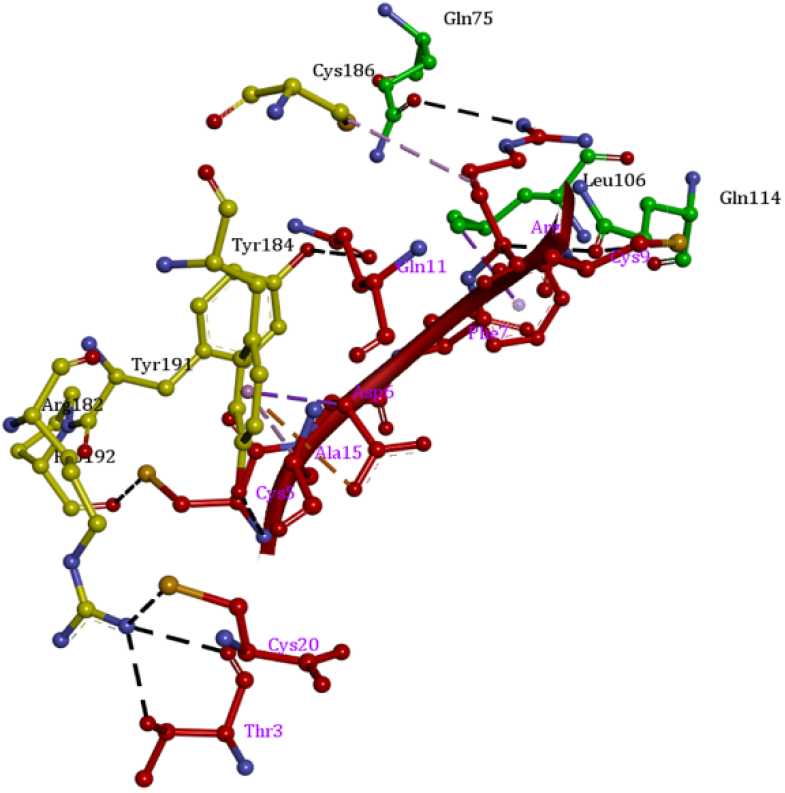
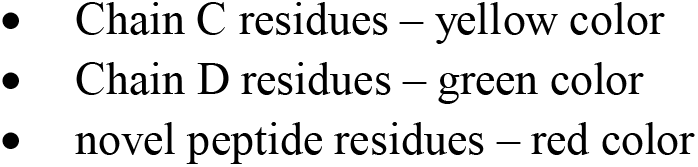
3D interaction details of novel (F8J2D7) peptide with nAChR pocket residues

The α**-BTx -peptide-nAChR** complex, exhibited 5 hydrogen bonds at chain C via the residues **THR195, ARG182, LYS141 and TYR91** along with 5 hydrophobic interactions across **ARG182 & TYR184**, and 3 electrostatic interaction from **ARG182 and ASP193** respectively. At chain D, 1 hydrogen bonds and 1 sulfur-pi-orbitals bonds were noted due to **GLY163 and TYR167** (Table 4). In case of, **novel peptide(F8J2D7)-nAChR complex**, 6 hydrogen bonds were found at chain C due to **ARG182, TYR191, TYR184, and PRO192**, along with 3 hydrophobic interaction from **TYR184 and CYS186** and 1 electrostatic interaction (negative-pi-orbitals type) due to **TYR184** respectively . At chain D, 5 hydrogen bonds and 1 hydrophobic interaction were observed due to **GLN157, SER162, GLN75** and **GLN114** (Table 4). On the other side, 1NTN-peptide-nAChR complex, showed only 1 hydrogen bond and 1 electrostatic interaction at chain C via the residues **TYR184 and ARG182**. At chain D, 4 hydrogen bonds and 2 hydrophobic interaction were noted due to **TYR164, GLY163, PRO166 and ILE165** respectively (Table 4).

Furthermore, the associated docking energy scores for the respective sets of peptide complexes, were estimated and noted down (Table 4). The results exhibited that, amongst the peptide systems, **the novel-peptide (F8J2D7)-nAChR complex** depicted the maximum docking energy score of **-255.17 kcal/mol** with 16 favorable interactions with nAChR pocket residues, followed by the α**-BTx -peptide-nAChR complex**, with an energy score of **-251.89 kcal/mol** with 15 favorable interactions respectively. On the other hand, the 1NTN-peptide complex, had the least docking energy score of **-173.59 kcal/mol,** with only 8 favorable number of interactions (refer Table 4). Thus, based on the results, we can conclude that both the peptide complexes i.e. **novel-peptide (F8J2D7)-nAChR complex** and the α**-BTx -peptide-nAChR complex**, exhibited strong favourable binding affinities with the receptor (nAChR), better than the complete toxin molecules.

### 3.4 Molecular Dynamics studies

The extent of binding affinities and structural stabilities of the docked complexes, were evaluated using Molecular Dynamics simulations studies **(Hollingsworth & Dror, 2018).**

#### 3.4.1 Root mean square deviation (RMSD) studies

The RMSD values for the complexes were evaluated using GROMACS command line gmx-rmsd using the trajectory information. The RMSD values indicate the stability of the complexes, and higher the value of the RMSD suggests, instability for the complexes. In our study, RMSD values (in Angstroms) for the respective complexes were computed for the c-alpha atoms and backbone atoms, respectively. Further, the maximum, minimum and average values were respectively tabulated, using the trajectory frames belonging to the last (25-70 ns) (Table 5). From the figures, we can conclude that, all the complexed structures initially showed upwards RMSD trend (Figure 6a-6e). We noticed that, the toxin complexes exhibited an unbalanced and fluctuating trajectories across the simulation runs, suggesting higher instability of these complexes (Figure 6a-6b). On the other hand, the peptide systems trajectories were found to converge at 25 ns, with a steadiest nature of RMSD tone across the entire 70 ns run (Figure 6c-6e). Amongst the peptide systems, the 1NTN-peptide-nAChR complex showed a higher RMSD profile than the other two peptide complexes. **(Figure 6d)**. However, the other two peptide complexes i.e., **novel-peptide (F8J2D7)-nAChR and** α**-BTx -peptide-nAChR** exhibited a RMSD fluctuation profile of lower than 3.0 A (Figure 6c,6e), suggesting that, these peptide structures were more balanced and stable throughout the simulation run.

**Figure 6a:**
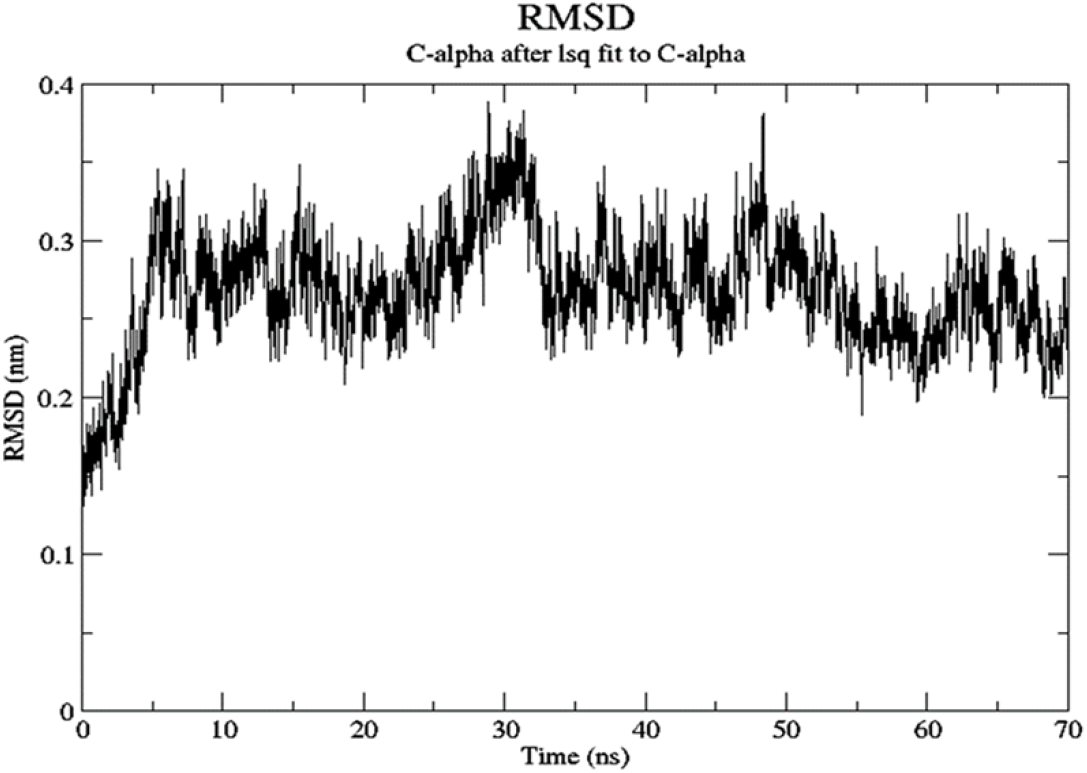
RMSD (c-alpha atoms) for α-BTx toxin-receptor complex;

**Figure 6b:**
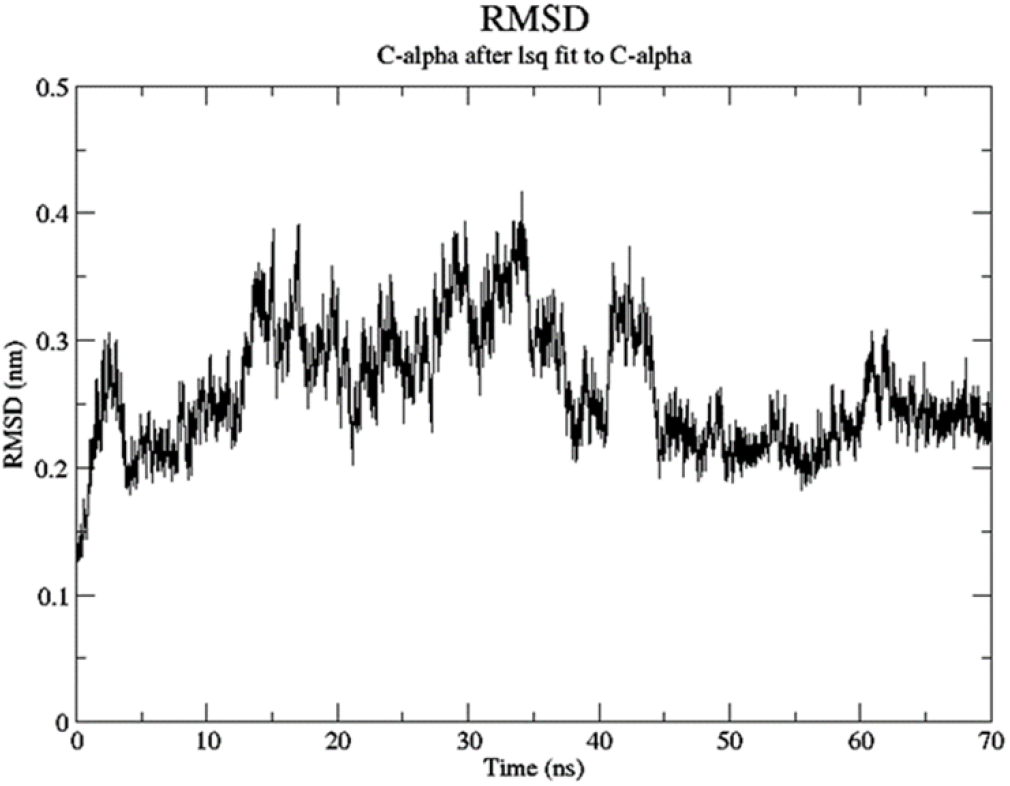
RMSD (c-alpha atoms) for 1NTN toxin-receptor complex

**Figure 6c:**
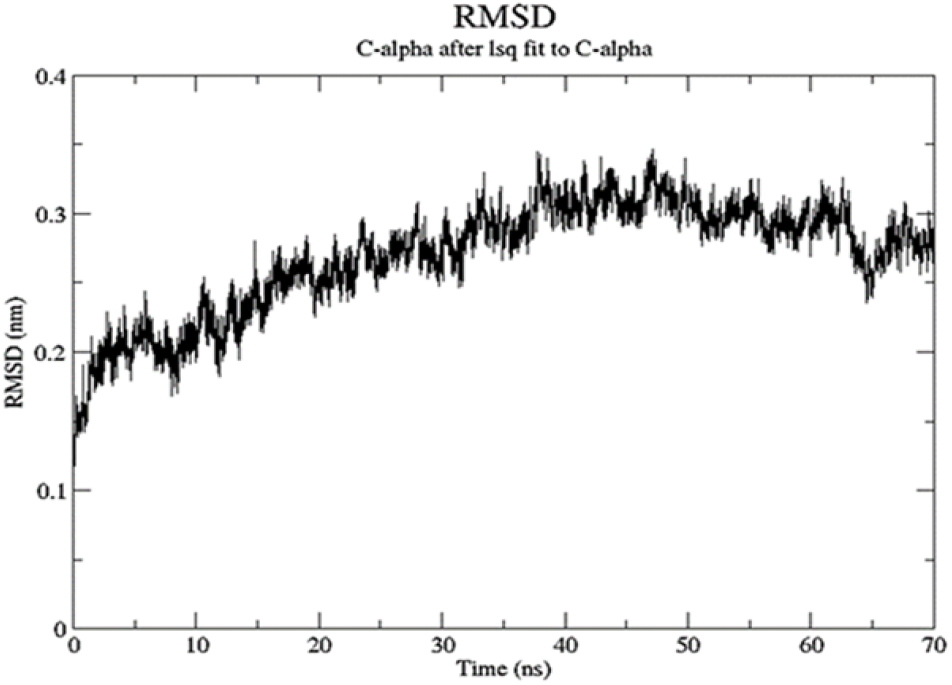
RMSD (c-alpha atoms) for novel peptide-receptor complex;

**Figure 6d:**
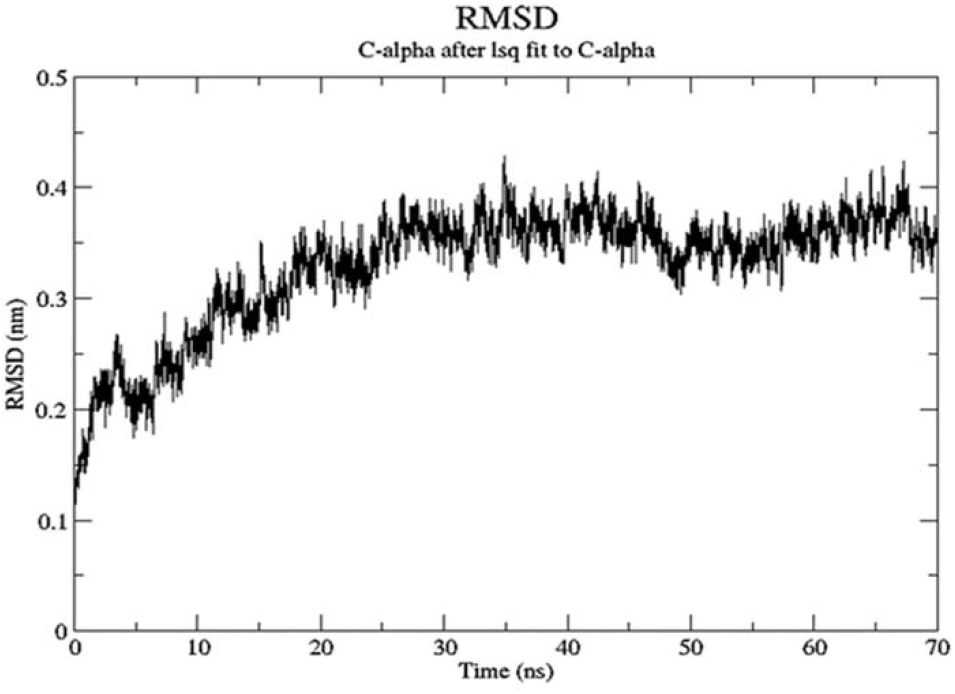
RMSD (c-alpha atoms) for INTN peptide-receptor complex

**Figure 6e:**
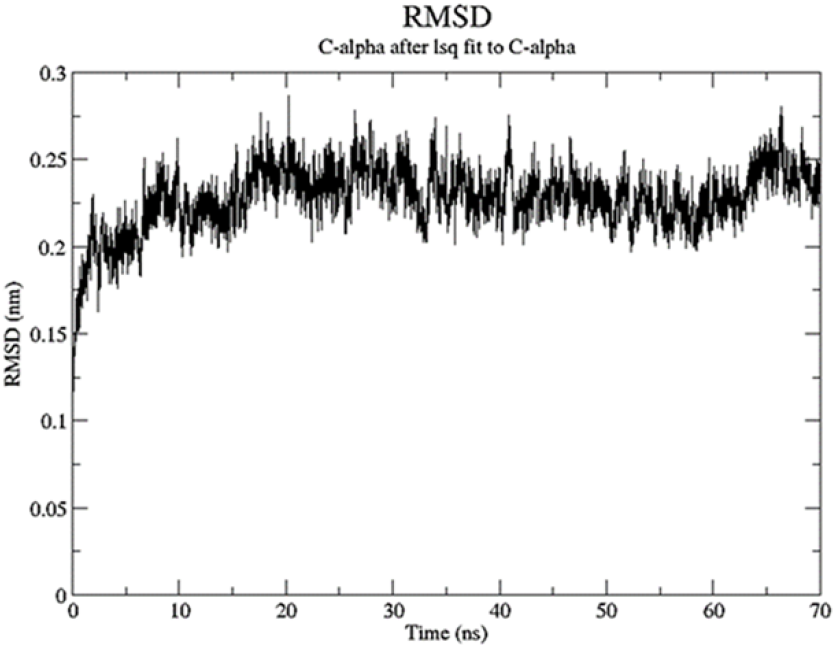
RMSD (c-alpha atoms) for alpha - BTx peptide-receptor complex

**Table 5:**
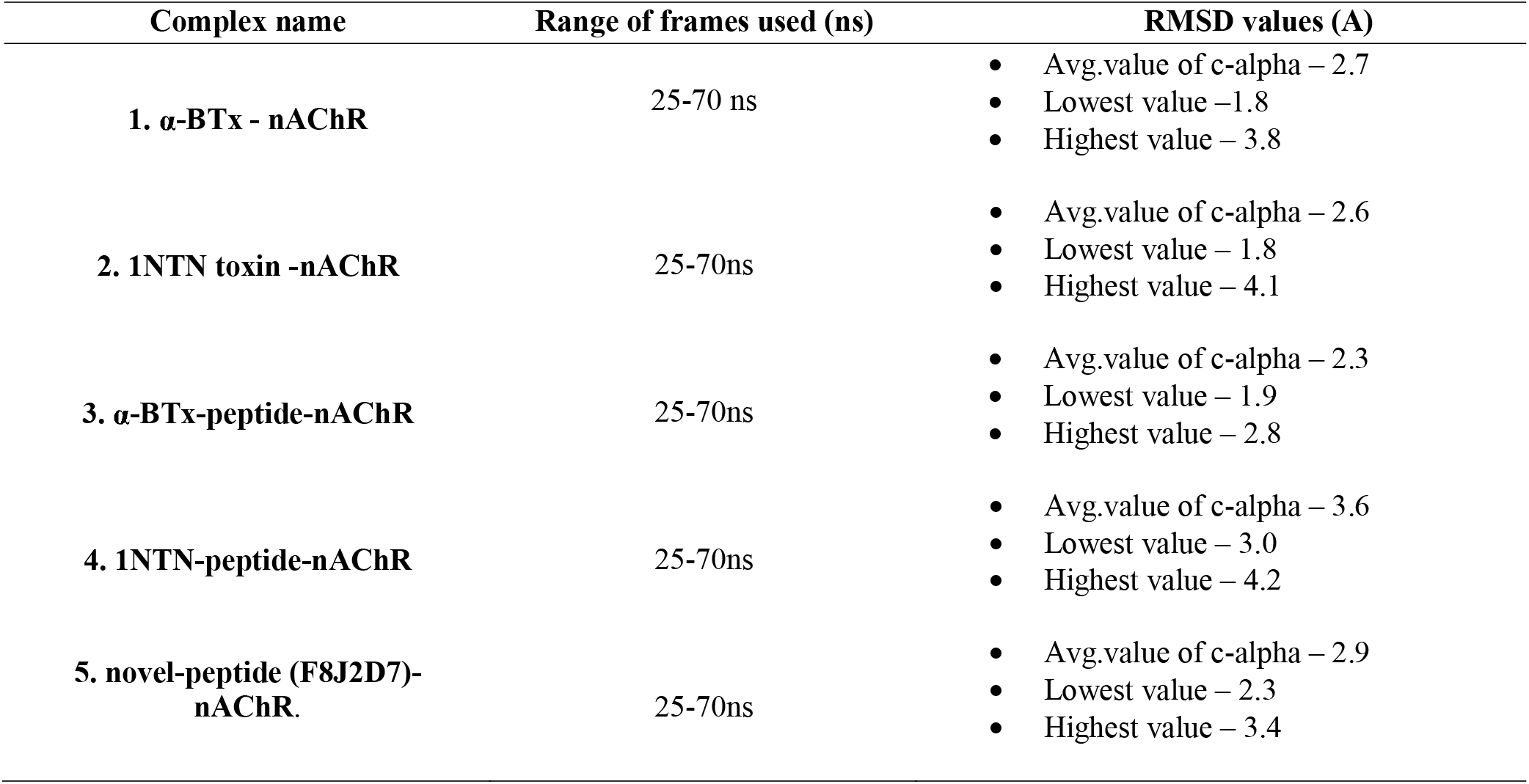
RMSD values of the complexes over 25-70ns trajectory frames

#### 3.4.2. Radius of Gyration (ROG) studies

To understand more about the stability of the complexes, the radius of gyration were computed using molecular trajectory information. The estimation of ROG values was done using GROMACS “gmx gyrate” command line. Radius of Gyration gives insights about the global level stability of either the ligand or the protein ternary structure, where gyration highlights the mass-weighted RMSD for a group of atoms in comparison to the common mass center. Hence, we can understand the stability and the compactness of the complexes of interest by depicting ROG values achieving a plateau around an average value. With our current study, we were able to confirm the stability and compactness of all of our complexes across the whole MD production run by the steadier ROG trajectories (Figure 7a-7e).Using the simulation trajectories of the complexes, we estimated the ROG values and, the average, maximum and the minimum gyrate values were also recorded and tabulated (Table 6). From the ROG profiles of the complexes, it was observed that, both the toxin structures i.e., α**-BTx - nAChR and 1NTN-nAChR** exhibited slightly higher average Rg values (nm) than the α-BTx peptide (2.36 nm) and novel peptide (2.37 nm) complexed systems (Table 6). Comparatively, a higher avg. Rg profile, was observed only for 1NTN-peptide-nAChR complex (2.43 nm) than other two peptide systems, indicating less rigidity of this complex. The other two peptide systems i.e., **novel-peptide(F8J2D7)-nAChR** and α**-BTx-peptide-nAChR** showed less deviations and had similar tone of Rg profiles throughout the 0-70 ns simulation run, which further validated the rigid nature of this peptide systems over the production run.

**Figure 7a:**
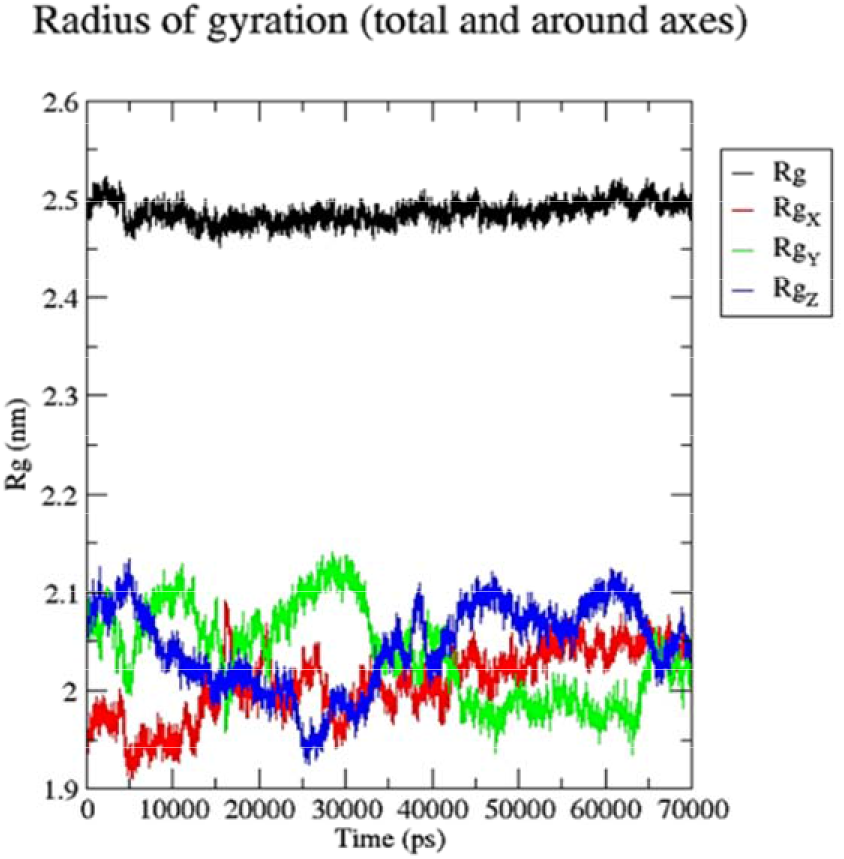
ROG fluctuation of α-BTx toxin-nAChR complex;

**Figure 7b:**
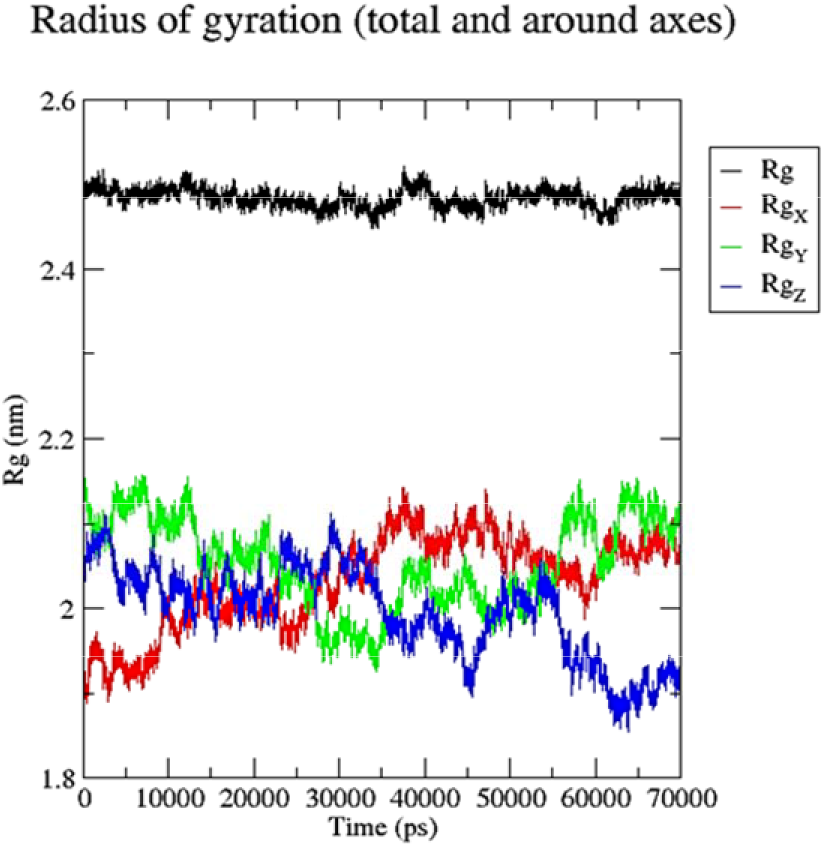
ROG fluctuation of 1NTN toxin -nAChR complex

**Figure 7c:**
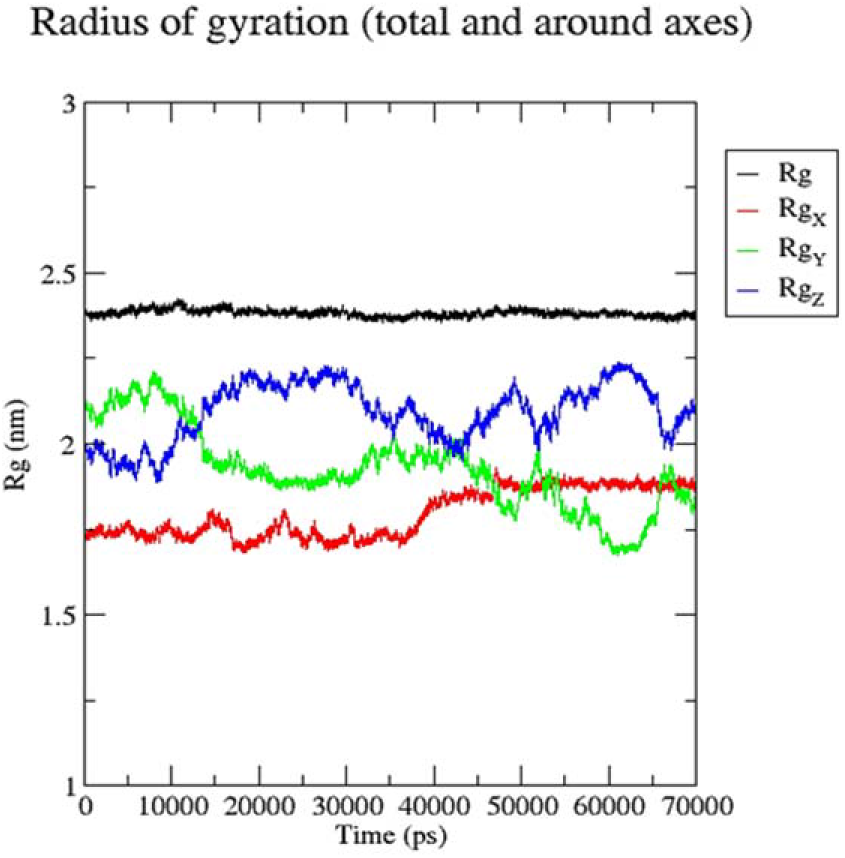
ROG fluctuation of novel peptide-nAChR complex;

**Figure 7d:**
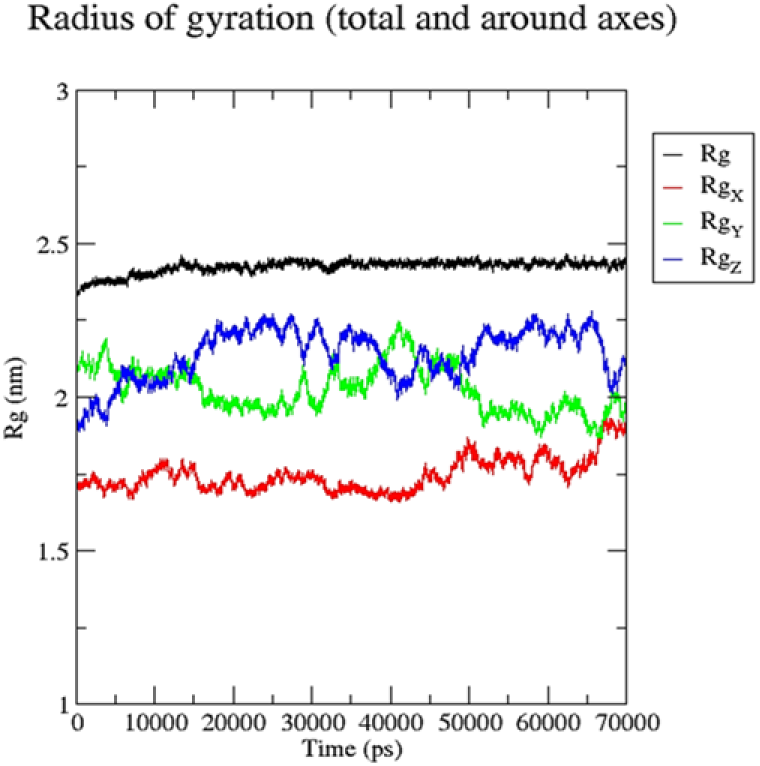
ROG fluctuation of 1NTN peptide-nAChR complex

**Figure 7e:**
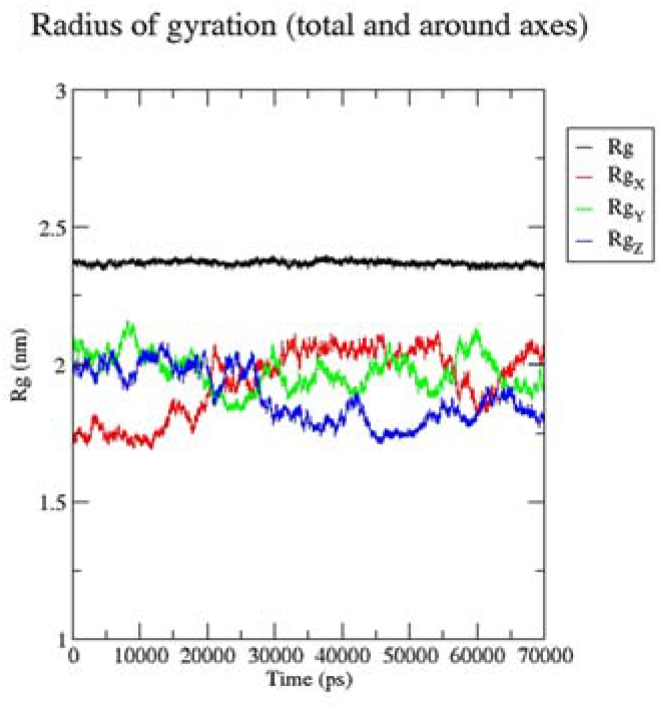
ROG fluctuation of α-BTx peptide – nAChR complex

**Table 6:**
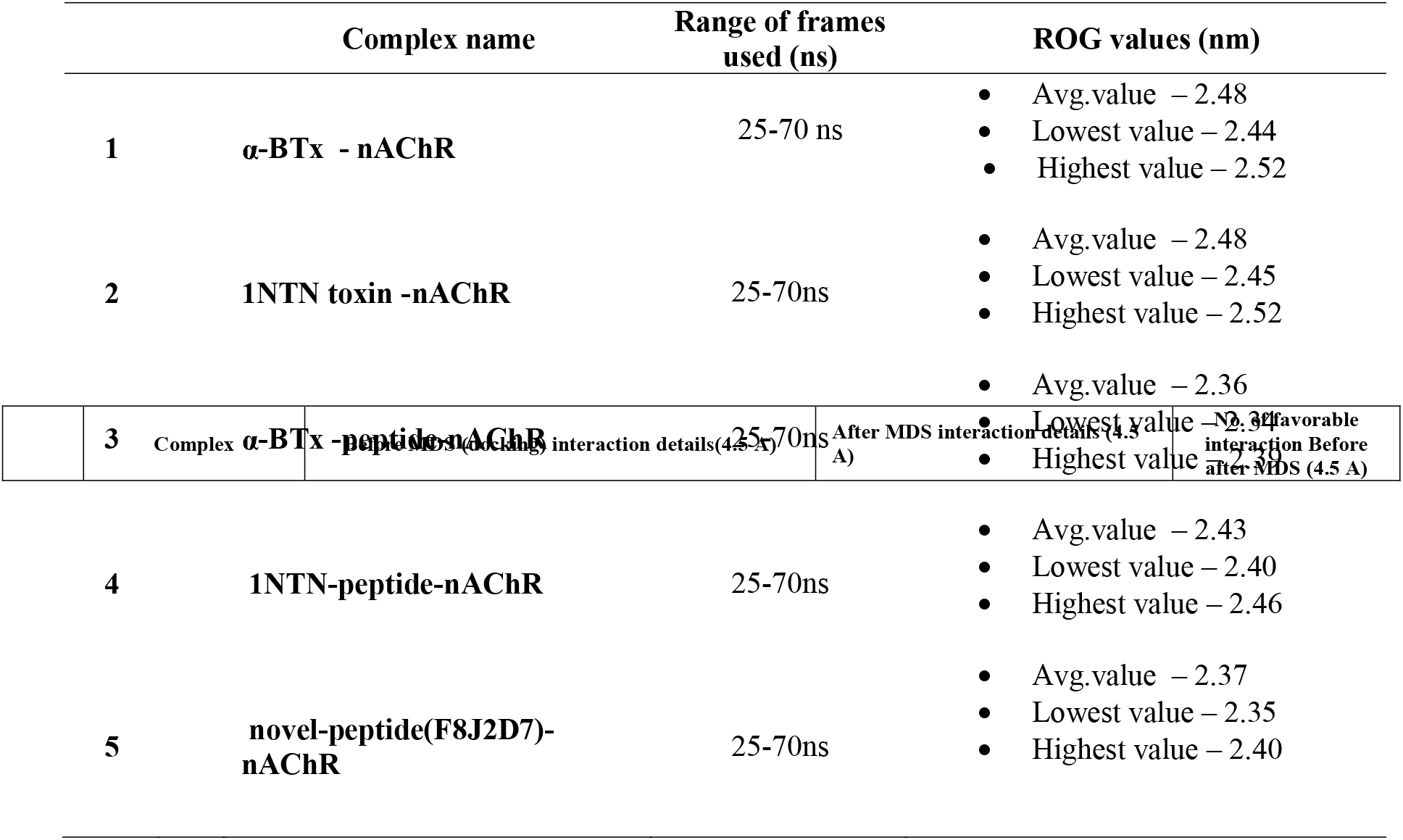
ROG values of the complexes over 25-70 ns trajectory frames

#### 3.4.3 Root mean square fluctuation (RMSF) and flexibility studies

In order to understand the flexible nature and the stability of the complexes in a dynamic environment, the per residue root-mean-square-fluctuation profile i.e. the average fluctuation of the atoms across the entire protein complex was evaluated for each complexed systems, using the GROMACS ‘gmx_rmsf’ command line. The RMSF value depicts the flexibility of the protein and it help us to understand and compute the time evolution of the average deviation for each residue from the referral position within the minimized starting structures. Likewise, using simulation trajectories of the complexes (25-70 ns), we had estimated the RMSF values and further the average, the maximum and the minimum rmsf values were also calculated and noted down. **(Table 7)**. From the RMSF profiles of the complexes, we noticed that, that the average atomic fluctuations tone was slightly higher for the bigger toxin structures **(Figure 8a-8b) (Table 7),** indicating less stability of this systems. However, amongst the protein-peptide systems, **the** α**-BTx peptide structure, comparatively** showed a lesser average atomic fluctuating profile (0.14 nm) than the other two peptide complexes i.e**., novel-peptide (F8J2D7)-nAChR** (0.15 nm) and 1NTN-peptide-nAChR complex (0.16 nm). Further, the figures **(Figure 8c, 8e**), indicating that the average fluctuating nature of almost every residue of this two peptide systems (α**-BTx peptide complex** and **novel-peptide complex)**, had RMSF atomic fluctuations lower than equal to 0.15 nm (1.5 A), suggesting the stability of this complexes over the simulation period.

**Figure 8a:**
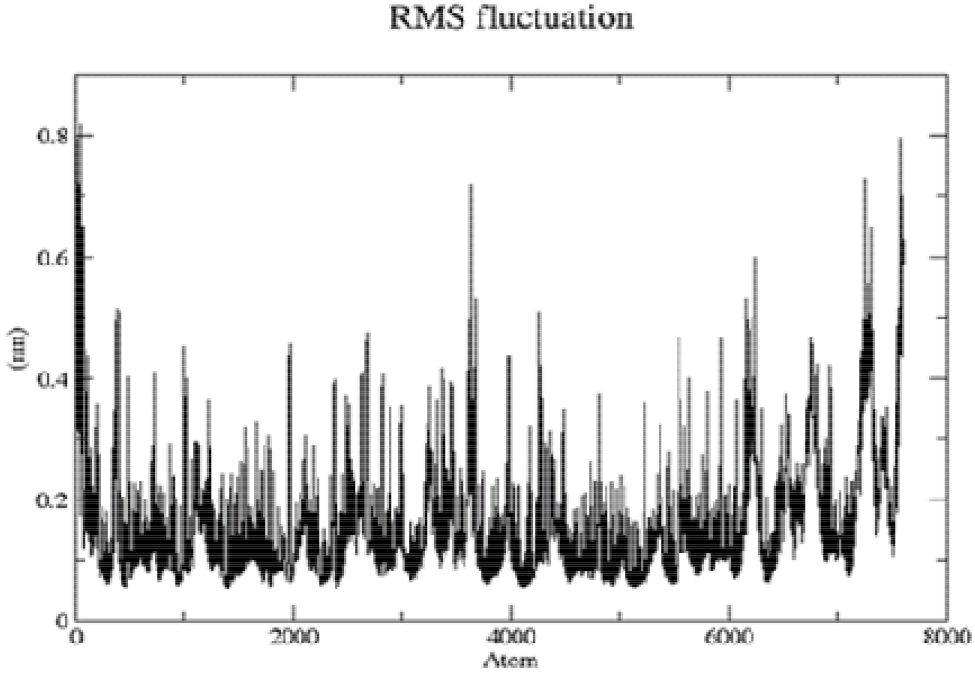
RMSF fluctuation profile of α-BTx toxin-nAChR complex;

**Figure 8b:**
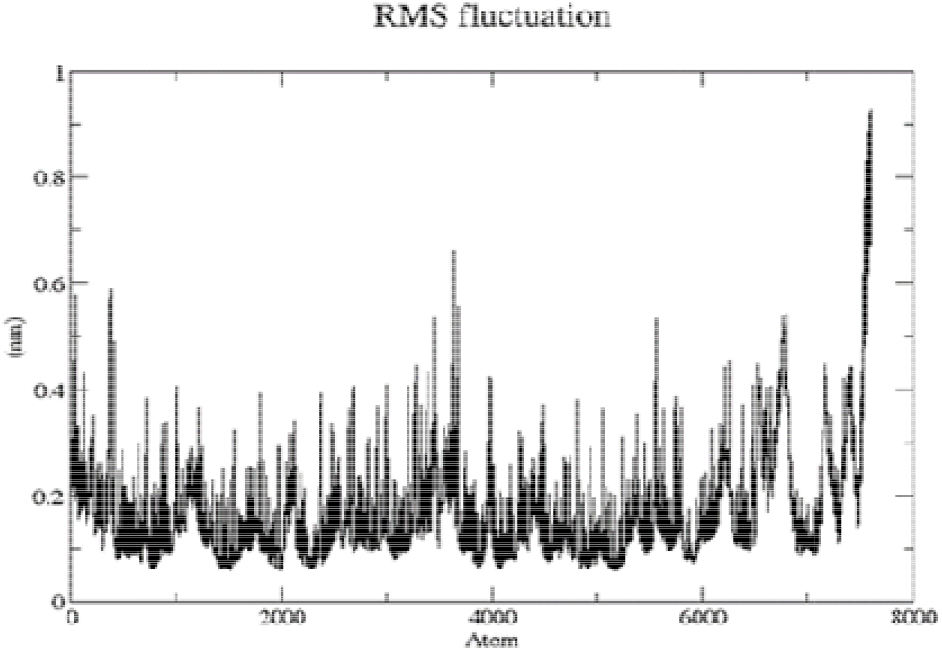
RMSF fluctuation profile of 1NTN -nAChR complex

**Figure 8c:**
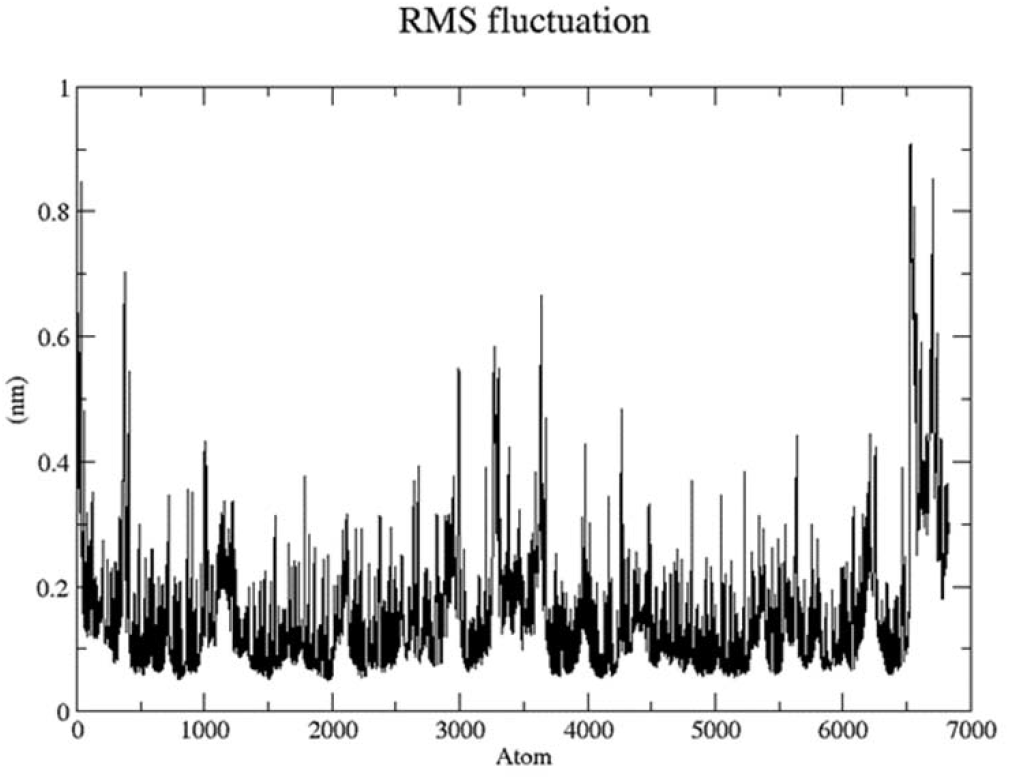
RMSF fluctuation profile of novel peptide-nAChR complex;

**Figure 8d:**
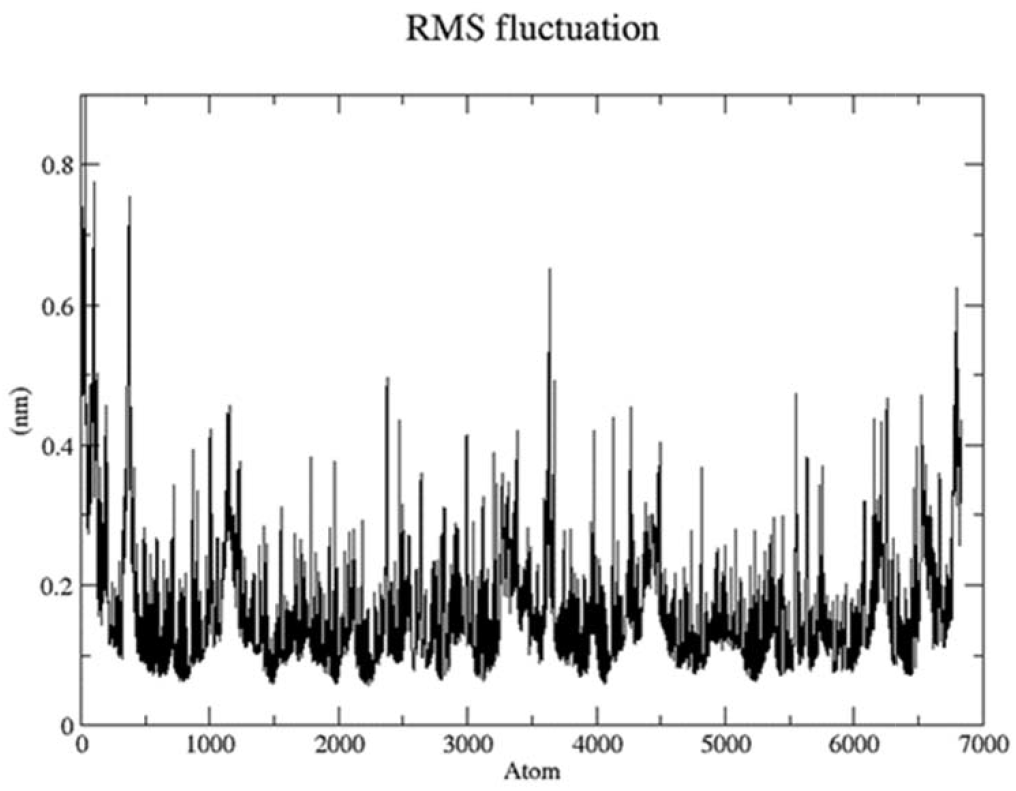
RMSF fluctuation profile of 1NTN peptide-nAChR complex

**Figure 8e:**
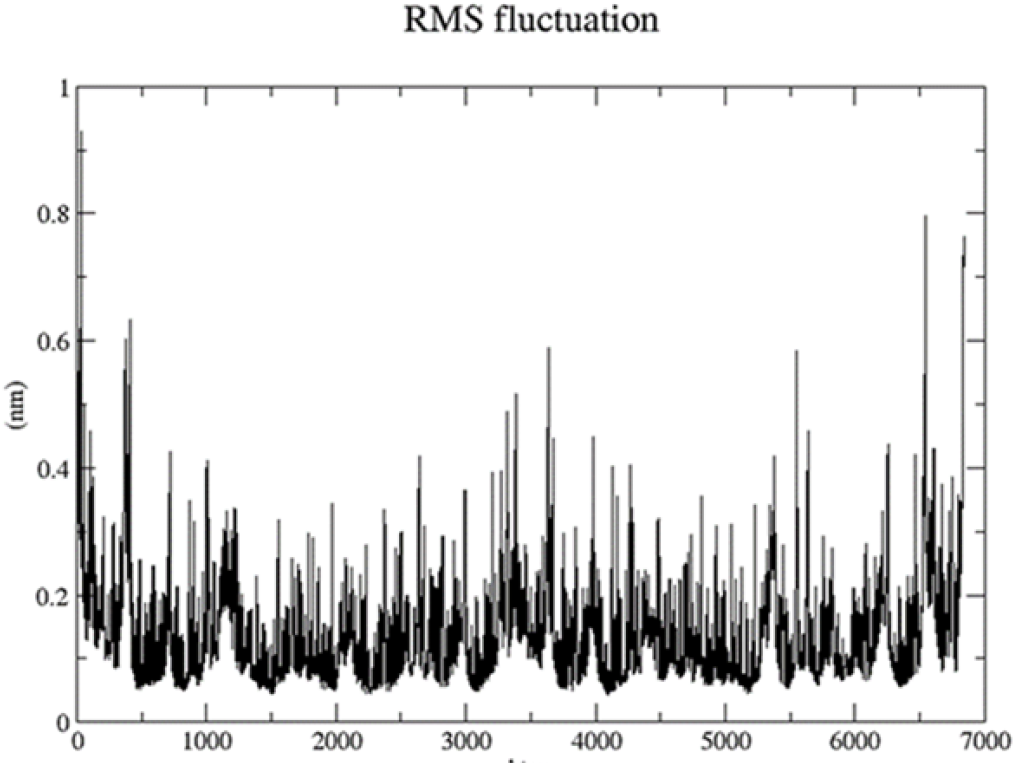
RMSF fluctuation profile for α-BTx peptide-nAChR complex

**Table 7:**
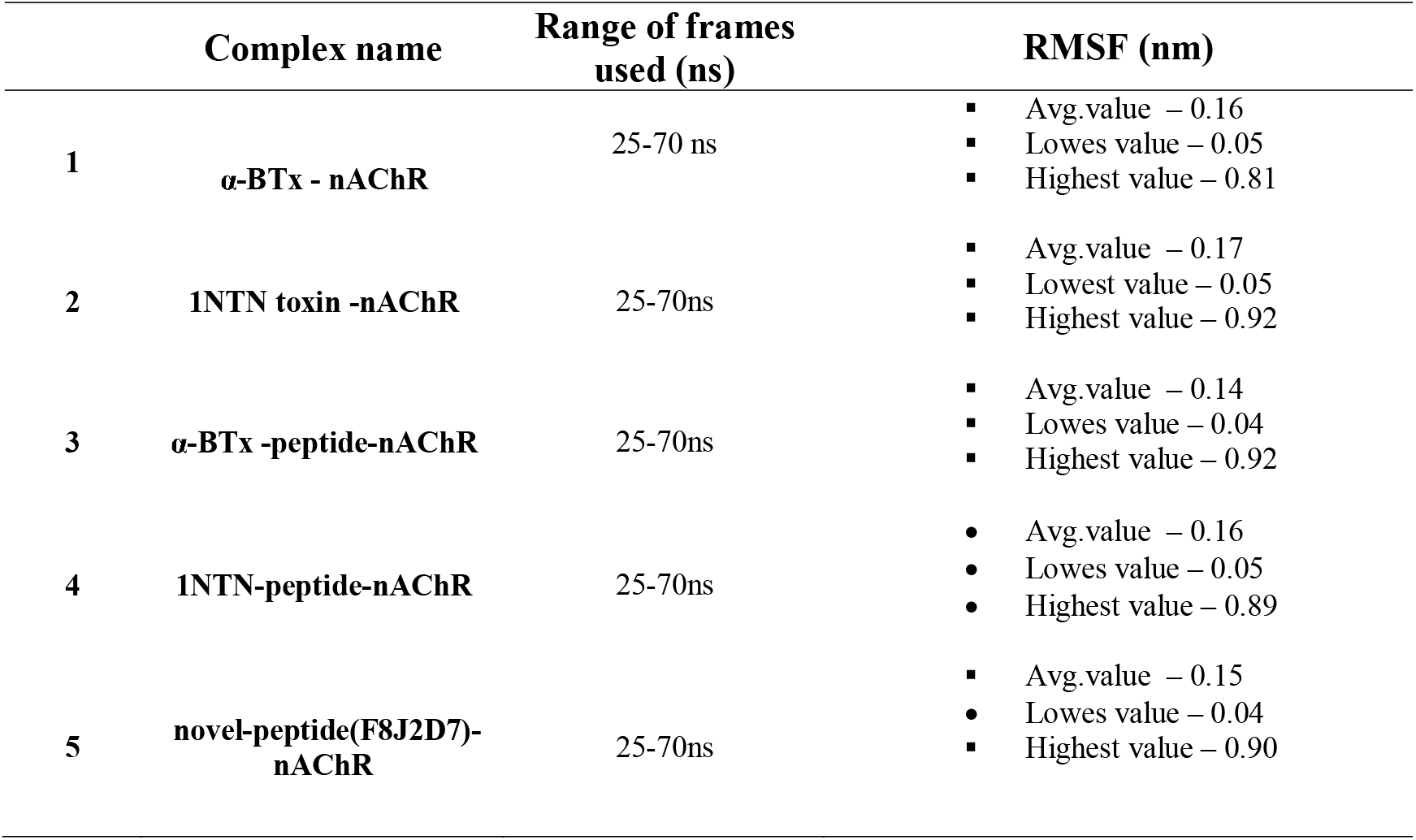
RMSF fluctuating values of the complexes over the trajectory frames (25-70 ns)

#### 3.4.4 Post Molecular dynamics interaction analysis

The simulation trajectories of the complexes, were used for understanding and analyzing the changes in the structural geometries as well as residual interaction, over the course of dynamic run. For this exercise, the simulation trajectories of the respective complexes were loaded in the VMD (visual molecular dynamics) tool **(Humphrey et al., 1996)**. and the coordinate frames information were extracted at different intervals across the whole 70 ns simulation run. Further, the retrieved coordinates of the complexes (at different intervals) were loaded in Discovery studio tool and likewise, the binding interaction analysis exercise (within 4.5 Angstroms) was carried out. The results from this analysis, were tabulated and compared with residual information gathered from initial docking exercise (Table 8).

**Table 8:**
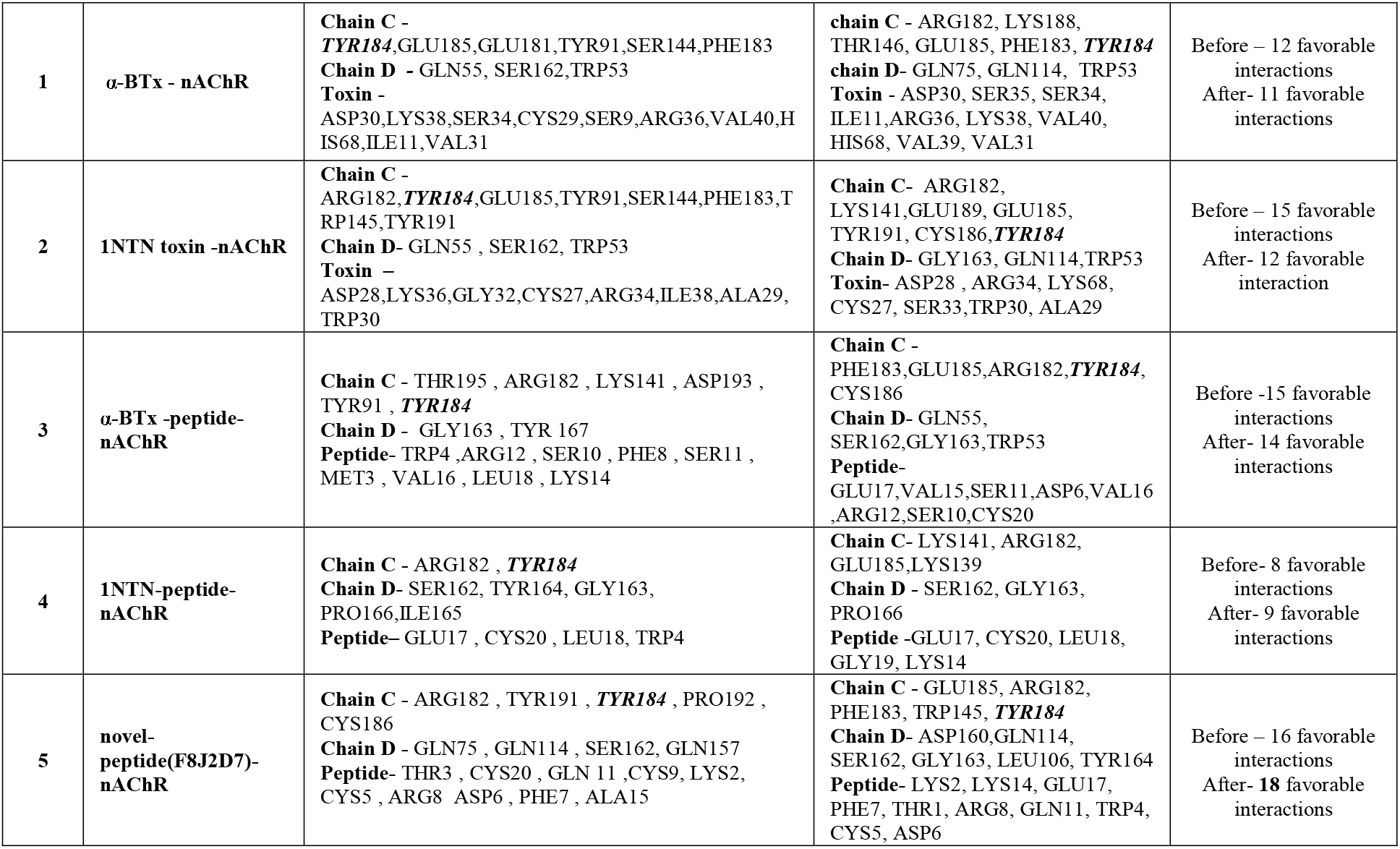
Details of interacting residues Before MD and post MD for the complexes

The post MD interaction analysis results exhibited that, amongst all the complexed systems, **the novel-peptide (F8J2D7)-nAChR complex** showed an increase in the molecular level interactions and comparatively had maximum number of favorable interactions (**18 favorable interactions**), followed by α**-BTx-peptide-nAChR complex (14 favorable interactions) (within 4.5 Angstroms**), suggesting stronger affinities of this peptides towards the receptor **(nAChR)** (Table 8). Further, the analysis results revealed that, all the complexed structures had maintained almost similar tone of binding chemistry, just like their initial docking interaction states (Table 8).

After 70 ns simulation run, we found that, the α**-BTx – nAChR toxin** complex had 11 favorable interactions, and was stabilized by 7 hydrogen bonds at chain C across the residues **ARG182, LYS188, THR146, GLU185, PHE183** and **TYR184**, along with 1 electrostatic interaction at **ARG182**. Further, at chain D, 2 hydrogen bonds and 1 hydrophobic interactions were noted due to **GLN75, GLN114 and TRP53** residues (Table 8). While in case of **1NTN toxin –nAChR complex,** 12 favorable interactions were noted and further this complex was found to be stabilized via 5 hydrogen bonds, along with 3 hydrophobic residues and 1 electrostatic interaction at chain C loop due to **ARG182,LYS141,GLU189,GLU185,TYR191,CYS186 and TYR184** residues (Table 8). Furthermore, 2 hydrogen bonds and 1 hydrophobic interaction were also noted at chain D via the residues **GLY163, GLN114 and TRP53** respectively (Table 8).

In case of peptide systems, after the simulation run, the α**-BTx -nAChR peptide** complex showed 14 favorable interactions and was stabilized by 6 hydrogen bonds at chain C via the residues **PHE183,GLU185,ARG182, and TYR184**, along with 1 hydrophobic interaction across **PHE183** and 1 other interaction (sulfur-O,N,S bond type) via **CYS186** respectively **(Table 8).** At chain D, 4 hydrogen bonds were noted due to **GLN55, SER162, and GLY163,** along with 2 hydrophobic interactions via **TRP53** . Interestingly, the **novel-(F8J2D7)-nAChR peptide complex**, showed **18 favorable interactions** and was found to be stabilized by 8 hydrogen bonds and 1 hydrophobic interaction at receptor chain C loop via the residues **GLU185, ARG182, PHE183, TRP145, and TYR184.** Again, at chain D, 7 hydrogen bonds and 2 hydrophobic interaction were noted due to **ASP160, GLN114, SER162, GLY163, LEU106 and TYR164 residues** respectively (Table 8).

Likewise, in case of **1NTN-peptide-nAChR complex**, we found that, this complex had 9 favorable interactions and after the simulation run, it was stabilized by 5 hydrogen bonds and 1 electrostatic interaction at chain C loop via **LYS141, GLU185, LYS139, ARG182** residues. At chain D, only 3 hydrogen bonds were noted due to **GLY163, PRO166,** and **SER162** residues (Table 8). Furthermore, we also noticed that, this particular peptide complex didn’t show any interaction with the critical amino acids like, **TYR184**, **PHE183** from chain C loop of the receptor (**nAChR**), suggesting less favorable affinity with receptor chain loops (Table 8).

During the simulation run of α**-BTx –nAChR** complex, the interactions at **ARG182, GLU185, PHE183, TYR184, GLN75, GLN114**, were found to be conserved and occupied throughout. Similarly, the interactions at **ARG182, GLU185, TYR191, CYS186, GLU189, GLY163, GLN114** for the 1NTN toxin-nAChR complex simulation.

Again, in case of α**-BTx-nAChR peptide** complex simulation, the interactions at residue **ARG182, PHE183, GLU 185, TYR184, GLN55 residues** were occupied throughout the simulation period, similarly, the interactions at residues **ARG182, GLU185, TRP145, ASP160, GLY163, GLN114, SER162** were found to be conserved and occupied throughout the whole novel-peptide(F8J2D7)-nAChR complex simulation run. But, in case of 1NTN-peptide complex, interaction only at residues **ARG182, SER162** were noticed throughout the simulations. Thus, from the post MD interaction analysis, we can conclude that, both the peptide systems, i.e., the α**-BTx-peptide-nAChR complex** and **novel-peptide(F8J2D7)-nAChR complex** showed good number of stronger favorable interactions with the receptor (nAChR) active site critical residues (chain C loop), suggesting that this two peptides may inhibit the nAChR with high binding affinities.

Furthermore, using the retrieved coordinate’s information from the simulation trajectories of the peptide systems, binding conformational analysis exercise at different intervals of ns across the pocket site of nAChR was carried out (Figure 9a-9c). From the figures, it was found that, comparatively, both the peptide systems (α**-BTx-peptide-nAChR complex** and **novel-peptide(F8J2D7)-nAChR complex**) showed more compacted geometries throughout the simulation run and these two small peptide moieties were found to be in a closed conformation states across the binding pocket area of nAChR., than 1NTN-peptide-nAChR complex system (Figure 9a-9c), indicating higher degree of binding affinities and stabilities of these peptide molecules across the binding pocket area of **nAChR** in dynamic environment.

**Figure 9a:**
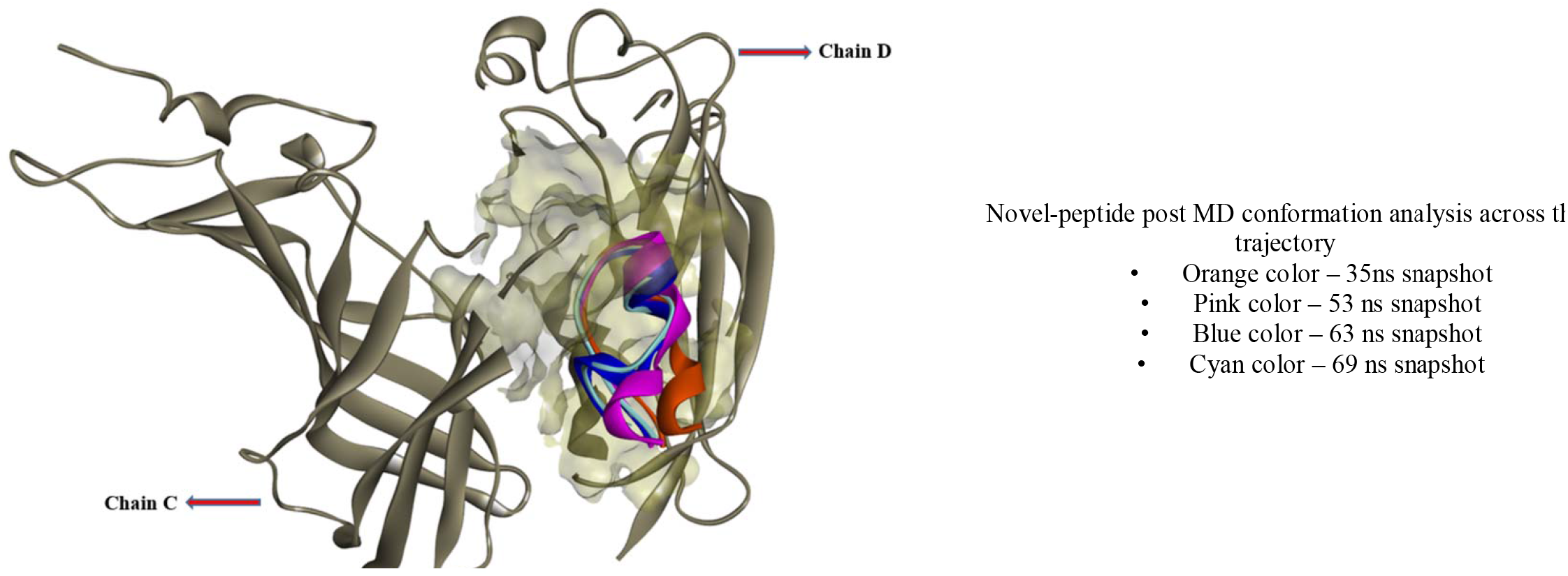
Binding nature of novel peptide inside the binding pocket area of nAChR (at different intervals)

**Figure 9b:**
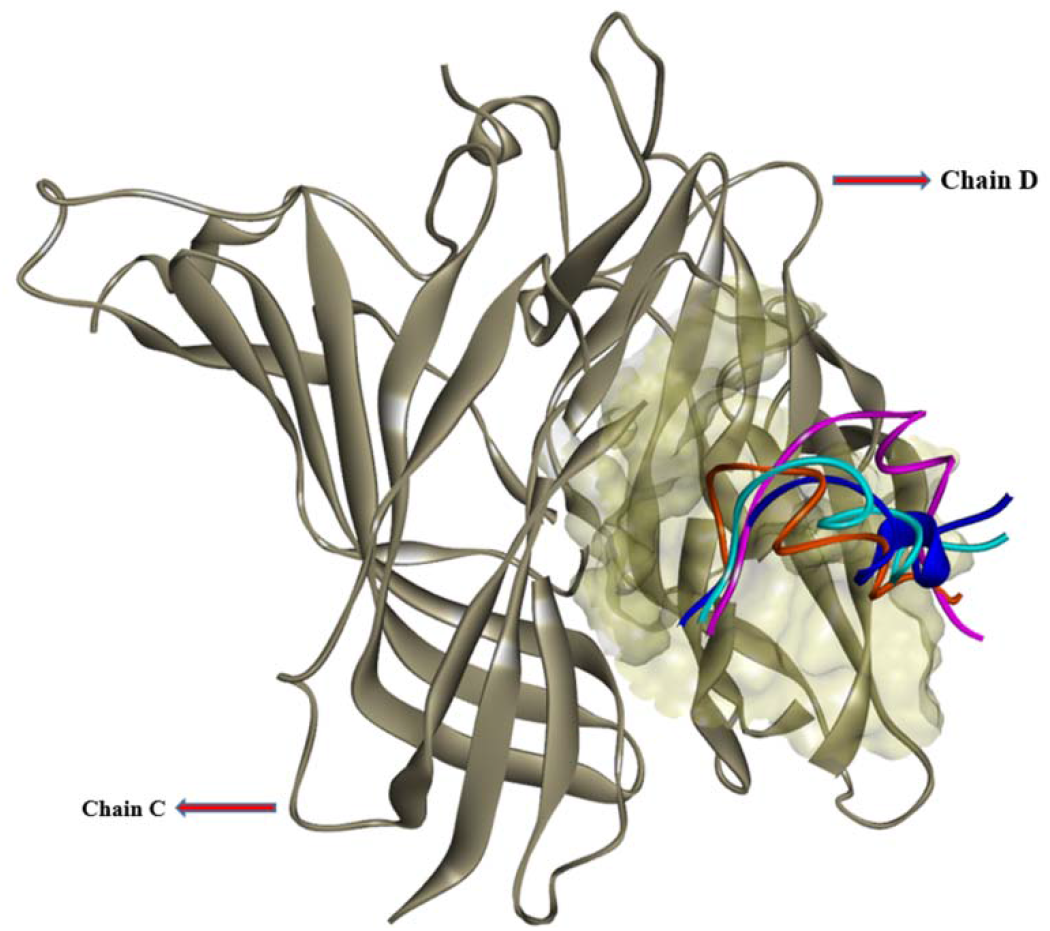
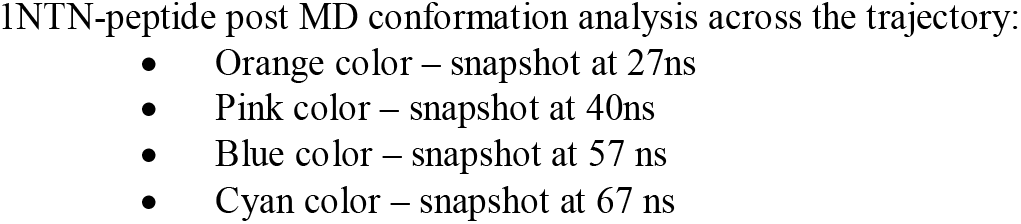
Binding nature of 1NTN peptide inside the binding pocket area of nAChR (at different intervals)

**Figure 9c:**
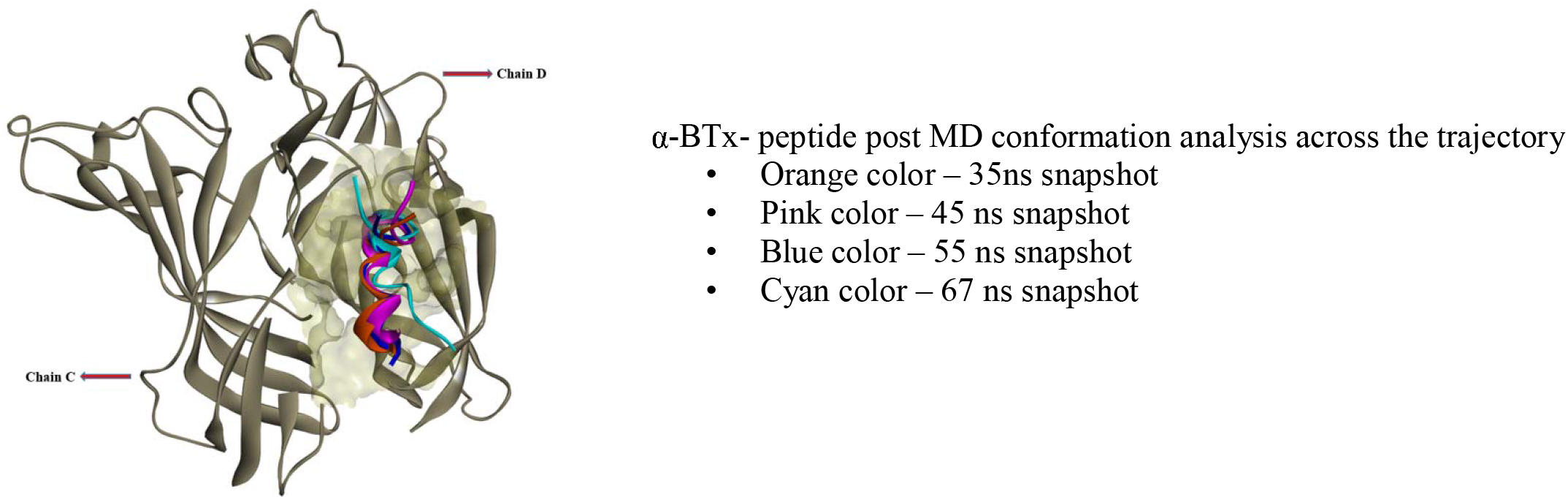
Binding nature of α-BTx- peptide inside the binding pocket area of nAChR (at different intervals)

#### 3.4.5 Average Binding free energy (MMPBSA calculation studies)

The binding free energy study was further carried out to revalidate the inhibitor affinity for the receptor – toxin, receptor-peptide complexes which was predicted by molecular docking studies. BFE calculation was carried out using Rashmi kumari MMPBSA module and was incorporated for the calculations using the “g_mmpbsa” tool on GROMACS. For this exercise, a total 100 ns frames for each of the complexes at an interval of 400 ps difference was extracted from the last 30-70ns trajectory and was used to calculate the BFE (kJ/mol) . The total average binding energy, MMPBSA energy components such as electrostatic, van der waals, polar solvation energy, non-polar solvation energy was calculated and tabulated below **(Table 9)**. The estimated results, exhibited that, amongst all the complexed systems, the **novel-peptide(F8J2D7)-nAChR peptide complex** had the maximum average total binding energy of **-1317.673 +/- 149.914 kJ/mol** followed by α**-BTx -peptide-nAChR complex** system with **-1145.516 +/- 143.978 kJ/mol (Table 9**), confirming stronger binding affinities of this two peptide systems than the complete toxins and the 1NTN peptide system. Furthermore, we noticed that, amongst the MMPBSA energy components, the van der Waals, electrostatic interactions, and the non-polar solvation energy had negatively contributed to the total binding energy, while the polar solvation energy component had positively contributed to total binding energy (**Table 9**). During the binding energy calculation study, the entropy impact factor over the complexes has not been considered due to higher computational cost.

**Table 9:**
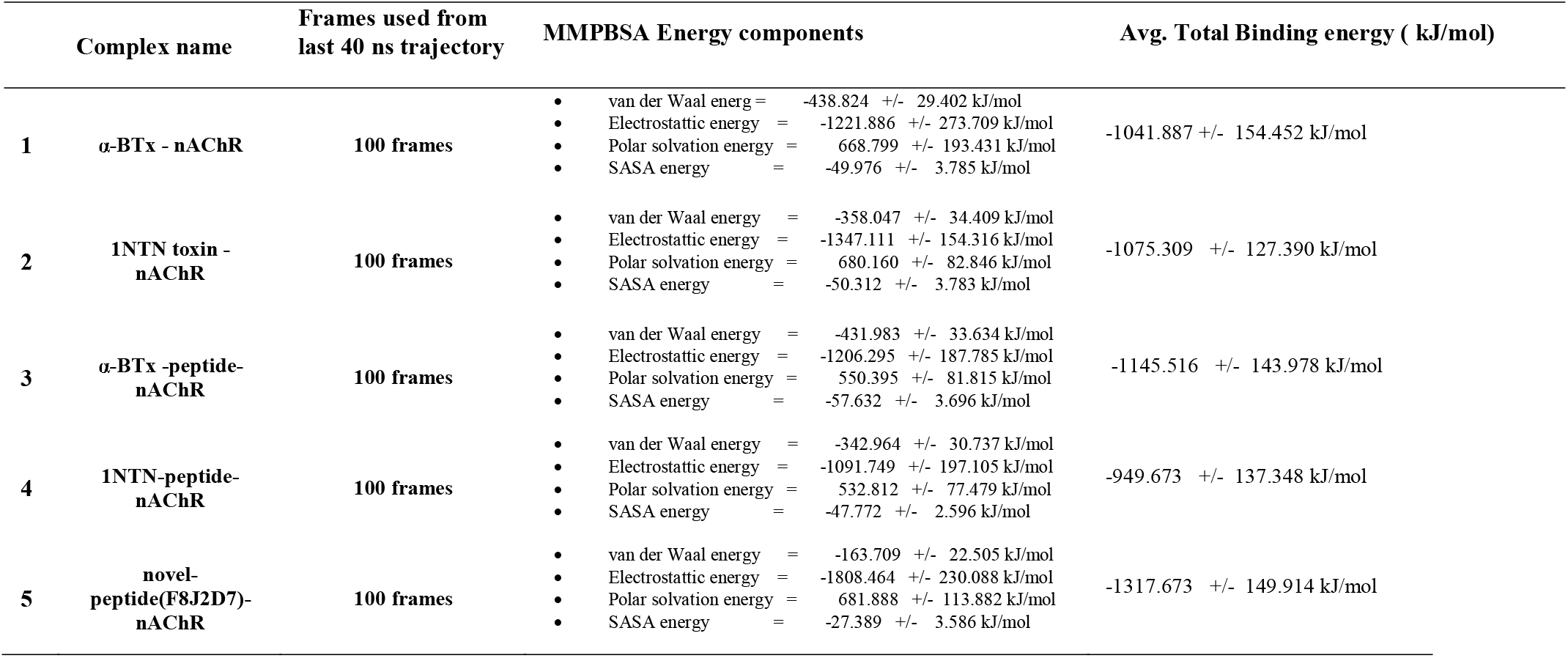
Binding-free calculations using MMPBSA module.

## Conclusion

To explore the development of novel therapeutics against severe involuntary muscle movement and trembling disorder (Dyskinesia), the study was conducted towards investigating the binding affinities of toxin proteins and short peptides, that could work as antagonistic leads for the nicotine acetylcholine receptor (nAChR) targets. The pharmacological interactions of snake venom toxins and its associated peptides with nAChR’s exhibit a promise towards exploiting these toxins as probable drugs for PD. Molecular docking and Molecular dynamics exercises were carried out using long chain neurotoxins (α-BTx, 1NTN) and the nAChR, with both the entire toxin and short peptide moieties. The study demonstrated that the novel-peptide(F8J2D7)- nAChR peptide complex had the maximum average binding energy of -1317.673 +/- 149.914 kJ/mol followed by α-BTx -peptide-nAChR complex system with -1145.516 +/- 143.978 kJ/mol, and hence these peptides appear to possess better affinity with the nAChR, which is a first step towards designing peptide-based drugs from snake venom proteins, that could facilitate further research in the peptide-drugs for various neurodegenerative diseases.

## Supporting information

Supplemental Interaction

## ABBREVIATIONS

1. nAChR: - Nicotine Acetylcholine Receptor
2. PD: - Parkinson Disease
3. NDD: - Neurodegenerative Diseases
4. mAChR: - Muscarinic Acetylcholine Receptor
5. GPCR: – G Protein Coupled Receptor
6. CNS: - Central Nervous System
7. LBD: - Ligand Binding Domain
8. MSA: - Multiple Sequence Alignment
9. PDB: – Protein Data Bank
10. FFT: - Fast Fourier Transform
11. BLAST: - Basic Local Alignment Search Tool
12. GROMACS: - GROningen MAchine for Chemical Simulations
13. MD: – Molecular Dynamics
14. RMSD: – Root Mean Square Deviation
15. RMSF: – Root Mean Square Fluctuation
16. ROG: – Radius Of Gyration
17. MMPBSA: - Molecular Mechanics Poisson-Boltzmann Surface Area

## Acknowledgements

The authors gratefully acknowledge the generous support and facilities extended by Sir M Visvesvaraya Institute of Technology and Sri Krishnadevaraya Educational Trust, Bangalore, towards this project.

