## Supplemental PDF for "Molecular Dynamics studies of Long Neurotoxins against nAChR provide key insights towards design of therapeutic peptides for Dyskinesia"

NCBI Multiple Sequence Alignment Viewer, Version 1.21.0

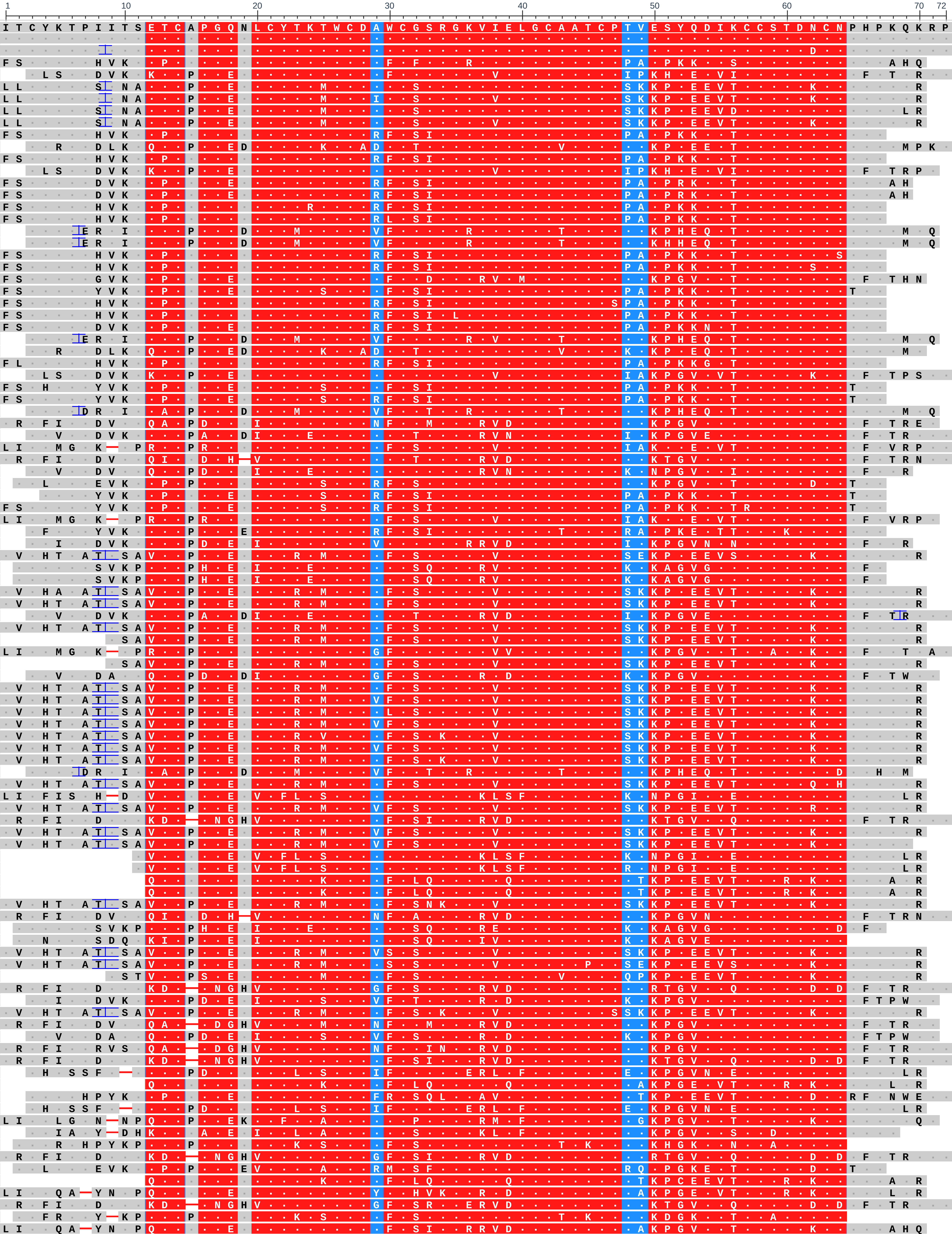


| Sequence ID | | Start | End | Organism |
|---|---|---|---|---|
| Query_16439 | (+) | 1 | 72 | |
| 1NTN_A | (+) | 1 | 72 | Naja oxiana |
| P01382.1 | (+) | 1 | 73 | Naja oxiana |
| A8HDK4.1 | (+) | 22 | 91 | Tropidechis carinatus |
| AHZ08823.1 | (+) | 24 | 93 | Micropechis ikaheca |
| A1IVR8.1 | (+) | 3 | 73 | Bungarus candidus |
| A1IVR9.1 | (+) | 3 | 73 | Bungarus candidus |
| C5ILC5.1 | (+) | 22 | 92 | Ophiophagus hannah |
| A1IVR7.1 | (+) | 3 | 73 | Bungarus candidus |
| ABX58158.1 | (+) | 22 | 88 | Austrelaps labialis |
| P0DQQ1.1 | (+) | 3 | 72 | Naja melanoleuca |
| ABX58156.1 | (+) | 22 | 88 | Austrelaps labialis |
| AHZ08824.1 | (+) | 24 | 92 | Micropechis ikaheca |
| ABX58155.1 | (+) | 22 | 90 | Austrelaps labialis |
| A8S6A8.1 | (+) | 22 | 90 | Austrelaps superbus |
| ABX58161.1 | (+) | 22 | 88 | Austrelaps labialis |
| ABX58163.1 | (+) | 22 | 88 | Austrelaps labialis |
| Q2VBP8.1 | (+) | 24 | 93 | Ophiophagus hannah |
| Q2VBP3.1 | (+) | 24 | 93 | Ophiophagus hannah |
| ABX58159.1 | (+) | 22 | 88 | Austrelaps labialis |
| ABX58162.1 | (+) | 22 | 88 | Austrelaps labialis |
| ABX58167.1 | (+) | 22 | 91 | Austrelaps labialis |
| F8J2E6.1 | (+) | 22 | 88 | Drysdalia coronoides |
| ABX58157.1 | (+) | 22 | 88 | Austrelaps labialis |
| ABX58160.1 | (+) | 22 | 88 | Austrelaps labialis |
| ABX58154.1 | (+) | 22 | 88 | Austrelaps labialis |
| Q2VBP6.1 | (+) | 24 | 93 | Ophiophagus hannah |
| P01383.1 | (+) | 3 | 70 | Naja melanoleuca |
| ABX58164.1 | (+) | 22 | 88 | Austrelaps labialis |
| AHZ08825.1 | (+) | 24 | 93 | Micropechis ikaheca |
| ACR78491.1 | (+) | 22 | 88 | Drysdalia coronoides |
| F8J2F2.1 | (+) | 22 | 88 | Drysdalia coronoides |
| P80156.1 | (+) | 3 | 72 | Ophiophagus hannah |
| P01389.1 | (+) | 1 | 71 | Naja anchietae |
| Q53B58.1 | (+) | 24 | 93 | Ophiophagus hannah |
| P01384.2 | (+) | 22 | 92 | Notechis scutatus scutatus |
| P0DQQ2.1 | (+) | 1 | 71 | Naja melanoleuca |
| Q53B57.1 | (+) | 24 | 90 | Ophiophagus hannah |
| BAN66265.1 | (+) | 23 | 88 | Pseudechis rossignolii |
| ACR78483.1 | (+) | 25 | 88 | Drysdalia coronoides |
| ACR78488.1 | (+) | 22 | 88 | Drysdalia coronoides |
| 0512217A | (+) | 1 | 70 | Notechis scutatus |
| Q7T2I3.1 | (+) | 23 | 87 | Laticauda laticaudata |
| Q53B53.1 | (+) | 23 | 89 | Ophiophagus hannah |
| AAC83997.1 | (+) | 1 | 72 | Bungarus multicinctus |
| 763050B | (+) | 2 | 67 | Dendroaspis sp. |
| P01395.1 | (+) | 2 | 67 | Dendroaspis viridis |
| CAM11303.1 | (+) | 3 | 74 | Bungarus candidus |
| 1ABT_A | (+) | 1 | 72 | |
| P01387.1 | (+) | 3 | 73 | Ophiophagus hannah |
| ADN67593.1 | (+) | 3 | 74 | Bungarus multicinctus |
| 1HOY_A | (+) | 11 | 72 | Bungarus multicinctus |
| B2BRQ5.1 | (+) | 22 | 92 | Austrelaps labialis |
| CAM11306.1 | (+) | 13 | 74 | Bungarus candidus |
| P01386.1 | (+) | 3 | 71 | Ophiophagus hannah |
| P60615.1 | (+) | 22 | 93 | Bungarus multicinctus |
| CAB50692.1 | (+) | 22 | 93 | Bungarus multicinctus ... |
| CAM11308.1 | (+) | 3 | 74 | Bungarus multicinctus |
| 4HQP_F | (+) | 1 | 72 | Bungarus multicinctus |
| AAC83985.1 | (+) | 1 | 72 | Bungarus multicinctus |
| 1HC9_A | (+) | 1 | 72 | Bungarus multicinctus |
| AAC83982.1 | (+) | 1 | 72 | Bungarus multicinctus |
| P80965.1 | (+) | 3 | 70 | Ophiophagus hannah |
| AAC83990.1 | (+) | 1 | 72 | Bungarus multicinctus |
| Q53B59.1 | (+) | 22 | 90 | Ophiophagus hannah |
| AAD41805.1 | (+) | 23 | 94 | Bungarus multicinctus |
| P01391.1 | (+) | 1 | 71 | Naja kaouthia |
| P60616.1 | (+) | 22 | 93 | Bungarus multicinctus |
| 7KOO_F | (+) | 1 | 71 | Bungarus multicinctus |
| P82662.2 | (+) | 31 | 90 | Ophiophagus hannah |
| Q2VBP4.1 | (+) | 31 | 90 | Ophiophagus hannah |
| ACR78480.1 | (+) | 32 | 90 | Drysdalia coronoides |
| F8J2E5.1 | (+) | 32 | 90 | Drysdalia coronoides |
| AAC83986.1 | (+) | 1 | 72 | Bungarus multicinctus |
| P01388.1 | (+) | 1 | 71 | Naja melanoleuca |
| P01394.1 | (+) | 2 | 67 | Dendroaspis viridis |
| P01397.2 | (+) | 2 | 64 | Dendroaspis polylepis ... |
| AAC83995.1 | (+) | 1 | 72 | Bungarus multicinctus |
| AAC83996.1 | (+) | 1 | 72 | Bungarus multicinctus |
| D2N116.1 | (+) | 13 | 74 | Bungarus caeruleus |
| P25669.1 | (+) | 1 | 71 | Naja naja |
| Q53B54.1 | (+) | 3 | 71 | Ophiophagus hannah |
| AAC83987.1 | (+) | 1 | 72 | Bungarus multicinctus |
| P25674.1 | (+) | 1 | 70 | Naja haje haje |
| A8N285.1 | (+) | 24 | 92 | Ophiophagus hannah |
| P34074.2 | (+) | 1 | 71 | Naja annulata annulata |
| P25671.1 | (+) | 1 | 71 | Naja naja |
| ETE56610.1 | (+) | 5 | 71 | Ophiophagus hannah |
| ACR78487.1 | (+) | 32 | 90 | Drysdalia coronoides |
| F8J2D7.1 | (+) | 24 | 93 | Drysdalia coronoides |
| Q53B56.1 | (+) | 24 | 90 | Ophiophagus hannah |
| ACY68696.1 | (+) | 22 | 91 | Parasuta nigriceps |
| P25670.1 | (+) | 3 | 67 | Aspidelaps scutatus |
| A3FM53.1 | (+) | 23 | 85 | Hydrophis hardwickii |
| P25668.1 | (+) | 1 | 71 | Naja naja |
| A6MFK4.1 | (+) | 23 | 88 | Demansia vestigiata |
| ACR78478.1 | (+) | 32 | 90 | Drysdalia coronoides |
| F8J2E1.1 | (+) | 22 | 90 | Drysdalia coronoides |
| P25673.1 | (+) | 1 | 71 | Naja naja |
| Q8UW29.1 | (+) | 23 | 84 | Hydrophis hardwickii |
| ACR78490.1 | (+) | 22 | 90 | Drysdalia coronoides |
